# Modelling vortex interactions reveals distinct patterns of brain reconfiguration following psilocybin and escitalopram

**DOI:** 10.64898/2026.11.06.764626

**Authors:** Álvaro Acosta-Gordo, Paulina Clara Dagnino, Yonatan Sanz Perl, Robin L. Carhart-Harris, David Erritzoe, David J. Nutt, Morten L. Kringelbach, Gustavo Deco

## Abstract

Major depressive disorder is a leading cause of disability, yet how interventions reorganise brain dynamics remains poorly understood. Here, we investigated how psilocybin and escitalopram reshape brain dynamics in depression using pre- and post-treatment resting-state functional magnetic resonance imaging (fMRI) from a double-blind randomised controlled trial (NCT03429075). We compared two whole-brain modelling approaches: conventional signal-space modelling describing interactions between regional brain signals, and vortex-space modelling capturing interactions between evolving patterns of local synchronisation. This represents the first application of vortex-space modelling to a psychiatric disorder. Vortex space was more sensitive than signal space in most analyses, including distinguishing treatment-related brain states, relating baseline connectivity to clinical improvement and predicting response to escitalopram. Our results further revealed distinct patterns of reorganisation following the two interventions. Psilocybin produced widespread increases in vortex-space effective connectivity and reduced network modularity, whereas escitalopram showed no comparable changes. Despite these differences, greater increases in brain turbulence (i.e., spatiotemporal variability in local synchronisation) were associated with greater symptom reduction in both treatments, with a significant post-treatment increase in responders. These findings suggest that psilocybin and escitalopram may reorganise brain dynamics in markedly different ways and yet share dynamical features associated with clinical improvement. Overall, vortex-space modelling offers a promising framework for characterising brain dynamics in depression and its successful treatment.

**Author Summary:** Psilocybin and escitalopram can both reduce depressive symptoms, but they may do so through different changes in brain dynamics. We analysed brain scans collected before and after treatment using two whole-brain modelling approaches. Signal-space modelling describes interactions between signals from individual brain regions, whereas vortex-space modelling captures how local patterns of synchronisation evolve and interact. This is the first application of vortex-space modelling to a psychiatric disorder. Overall, vortex space captured treatment-related and clinically relevant differences more effectively than signal space. Psilocybin produced widespread increases in vortex-space effective connectivity and reduced network modularity, whereas escitalopram did not. Nevertheless, increased brain turbulence was associated with greater symptom improvement in both groups. These findings suggest that the two treatments may reorganise brain dynamics through distinct routes while sharing dynamical features related to clinical improvement.

## Introduction

Major depressive disorder (MDD) is one of the leading causes of disability worldwide and imposes a substantial personal, social, and economic burden (Mnookin, 2016; World Health Organization, 2025). Although antidepressants are effective (Cipriani et al., 2018), their clinical benefits often take several weeks to emerge, many patients do not achieve remission after initial treatment, and relapse remains common (Otte et al., 2016; Rush et al., 2006). Treatment response also varies considerably between individuals (Nunez et al., 2023), making it difficult to anticipate which patients will benefit from a given intervention. Understanding how effective treatments act on the brain is therefore a priority.

Escitalopram and psilocybin represent two distinct approaches to the treatment of depression. Escitalopram is a conventional first-line antidepressant belonging to the class of selective serotonin reuptake inhibitors (SSRIs). It blocks the serotonin transporter, thereby increasing extracellular serotonin (Landy et al., 2023; Lundberg et al., 2007). Psilocybin, by contrast, is a classic serotonergic psychedelic administered in a limited number of dosing sessions within a psychologically supported therapeutic context (Rucker, 2024). Its active metabolite, psilocin, acts primarily as a partial agonist at serotonin 2A (5-HT2A) receptors (Madsen et al., 2019; Nichols, 2016). Yet, the broader functional implications of these molecular mechanisms continue to be investigated. A central unresolved question is how each treatment reshapes large-scale brain dynamics and how this reorganisation relates to clinical improvement.

One promising approach to this question draws on an emerging field in neuroscience that uses fluid dynamics to characterise the spatiotemporal organisation of the brain. In fluids, turbulence is characterised by irregular, swirling motion, in which vortices (i.e., coherent regions of rotating fluid) of different sizes interact, enabling mixing and energy transfer across scales. Analogously, brain activity exhibits turbulent-like fluctuations in local phase synchronisation (Deco & Kringelbach, 2020), with power-law scaling consistent with efficient information transfer across scales (Deco et al., 2023; Sanz Perl, Mininni, et al., 2023). This regime is thought to balance the stability required to maintain coherent function with the flexibility needed to adapt to changing internal and external demands. Importantly, disruptions in this balance have been associated with pathological brain states (Deco et al., 2025; Escrichs et al., 2022). In MDD, lower baseline turbulence has been associated with poorer subsequent response to escitalopram (Escrichs et al., 2025), whereas acute psilocybin administration has been shown to increase brain turbulence in healthy individuals, particularly at longer spatial scales (Cruzat et al., 2022). Nevertheless, these measures of turbulence remain purely descriptive: they characterise how brain dynamics differ across states, but not the mechanisms that give rise to them.

Whole-brain modelling offers a generative framework for investigating the mechanisms underlying brain dynamics (Deco & Kringelbach, 2014). Brain regions are represented as coupled oscillators whose interactions are constrained by empirical structural connectivity. Simulated dynamics are fitted to empirical functional connectivity by adjusting the weights of existing anatomical connections, yielding generative effective connectivity (GEC), the model-derived coupling required to reproduce the observed functional organisation (Deco et al., 2017). This approach has been successfully applied across different brain states and conditions (Kringelbach et al., 2020; Patow et al., 2024) and has revealed contrasting effects of psilocybin and escitalopram on large-scale brain organisation. By incorporating temporal asymmetry, Deco et al. (2024) found that psilocybin reduced the directedness of the brain’s functional hierarchy, whereas escitalopram increased it. Using in silico perturbations within this framework, Socoró-Garrigosa et al. (2025) found increased brain susceptibility following psilocybin but reduced susceptibility following escitalopram. Both studies, however, fitted models directly to regional blood-oxygen-level-dependent (BOLD) signals (i.e., in signal space), potentially missing informative signatures of turbulence previously observed in the brain.

Recent work has extended whole-brain modelling to a mesoscopic representation that explicitly accounts for these signatures by modelling the interactions between turbulent vortices (Deco et al., 2026). In the resulting vortex space, each brain region is represented through a local Kuramoto order parameter capturing the evolving amplitude and phase of synchronisation among its constituent signals. The whole-brain model is fitted to empirical functional connectivity between local synchronisation patterns, yielding generative effective connectivity in vortex space (GECVO), which describes the coupling required to reproduce the observed interactions. By incorporating this spatiotemporal organisation, vortex-space modelling may reveal features of brain dynamics that are not fully captured in signal space.

Here, we apply vortex-space whole-brain modelling to a psychiatric disorder for the first time. Resting-state fMRI and clinical data were acquired before and after treatment in a double-blind phase II randomised controlled trial comparing psilocybin and escitalopram in patients with MDD (Carhart-Harris et al., 2021). For each participant and session, we built signal- and vortex-space Hopf whole-brain models to compare their sensitivity to treatment-related brain reconfiguration. We further examined the clinical relevance of model-derived brain connectivity and empirical turbulence. We found that psilocybin and escitalopram produced distinct patterns of brain reorganisation, with vortex space being more sensitive to these changes than signal space in most analyses. Overall, vortex-space whole-brain modelling offers a principled framework for characterising treatment-related changes in brain dynamics and their clinical relevance.

## Results

In this study, we investigated how psilocybin and escitalopram reshape large-scale brain dynamics in MDD and whether treatment-related reconfiguration is captured differently in vortex and signal spaces. As shown in **Figure 1A**, resting-state fMRI data were acquired before and after treatment in a double-blind phase II randomised controlled trial comparing the two interventions (Carhart-Harris et al., 2021; details provided in *Methods*). We analysed these data using the recently introduced vortex-space whole-brain modelling framework (Deco et al., 2026) alongside conventional signal-space modelling (Deco et al., 2017).

**Figure 1.**
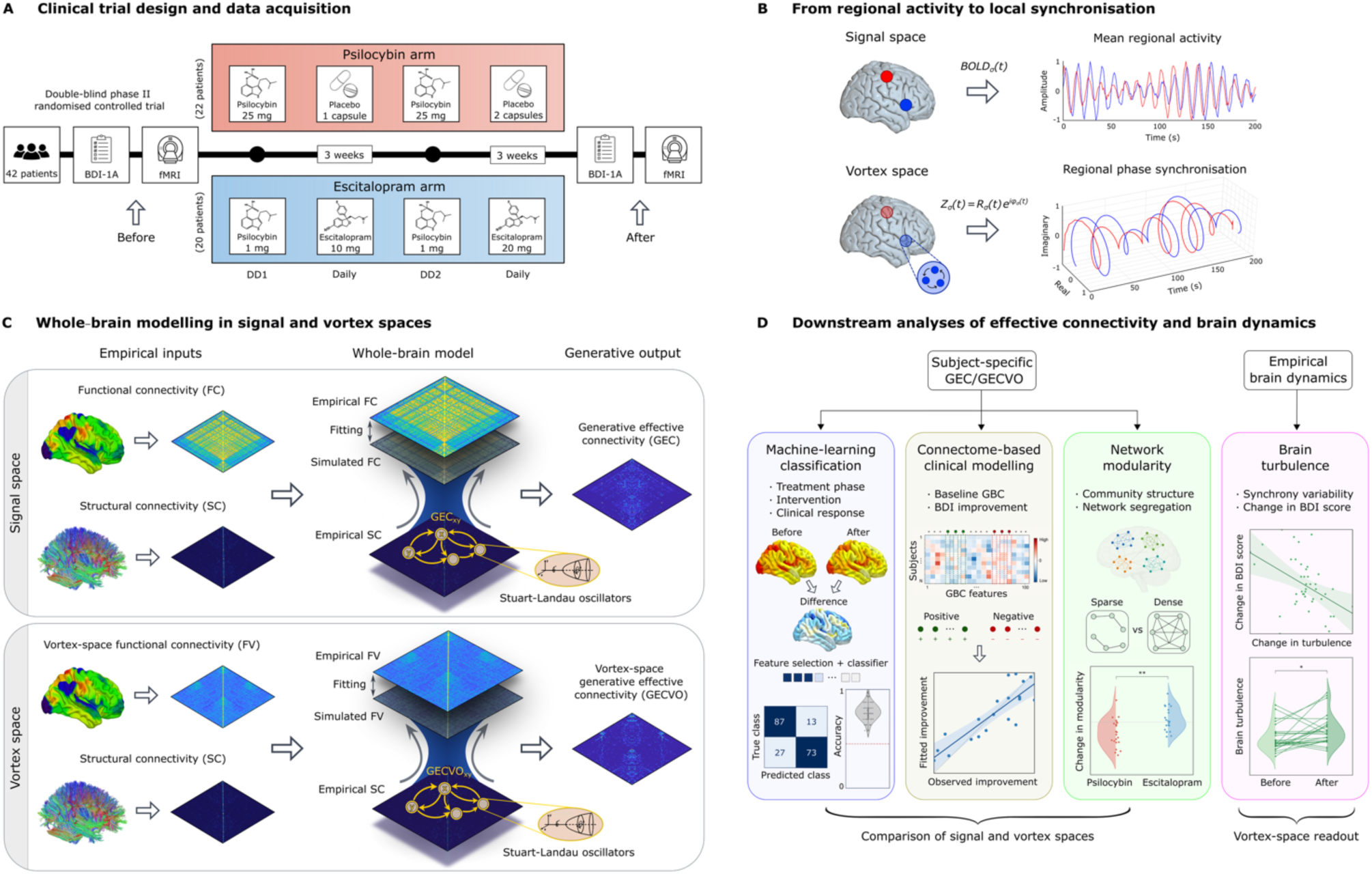
Conceptual framework for assessing treatment-related reconfiguration of brain dynamics across signal and vortex spaces in major depressive disorder. **A.** Resting-state neuroimaging data and clinical assessments were acquired before and after treatment within a double-blind phase II randomised controlled trial comparing psilocybin and escitalopram in patients with MDD. The panel shows participant allocation, the acquisition schedule, and the treatment regimens and doses administered in each arm. B. Signal and vortex spaces were defined over the same Schaefer-100 macroregions. In signal space, each macroregion was characterised by the mean BOLD activity of its constituent Schaefer-1000 parcels; in vortex space, it was characterised by local phase synchronisation among the same signals, quantified using a local Kuramoto order parameter. C. Whole-brain models were built for each participant and session independently in signal and vortex spaces. Empirical functional connectivity in signal space (FC) or vortex space (FV), together with structural connectivity (SC), was used to fit Hopf whole-brain models composed of coupled Stuart–Landau oscillators. Model fitting iteratively adjusted the weights of the existing structural connections to reproduce the corresponding empirical connectivity, yielding generative effective connectivity in signal space (GEC) or vortex space (GECVO). D. Subject-specific GEC and GECVO matrices were used for machine-learning classification of treatment phase and intervention, and prediction of clinical response from baseline connectivity; connectome-based clinical modelling relating baseline global brain connectivity (GBC) to subsequent symptom improvement; and network modularity analyses of treatment-related reconfiguration. These analyses enabled direct comparison between signal and vortex spaces. In parallel, empirical vortex-space dynamics were used to quantify brain turbulence and assess its relationship with clinical improvement.

### Whole-brain modelling in signal and vortex spaces

To compare the two representations at a matched anatomical scale, both were derived from the same BOLD time series parcellated using the Schaefer-1000 atlas and expressed over the same 100 macroregions of the Schaefer-100 parcellation (Schaefer et al., 2018). In signal space, each macroregion was characterised by the mean BOLD signal of its constituent fine-grained parcels; in vortex space, it was characterised by the local phase synchronisation among the same constituent signals, quantified using a local Kuramoto order parameter (**Figure 1B**). For each participant and session, we then constructed whole-brain models in both spaces using coupled Stuart–Landau oscillators constrained by the same empirical structural connectivity (**Figure 1C)**. The signal-space model was fitted to conventional functional connectivity (FC) between regional BOLD time series, yielding generative effective connectivity (GEC), whereas the vortex-space model was fitted to functional connectivity between local synchronisation patterns (FV), yielding generative effective connectivity in vortex space (GECVO).

The two implementations thus differed systematically in the empirical representation used for model fitting—regional BOLD activity in signal space and local synchronisation dynamics in vortex space—while retaining a common generative architecture (see *Methods*). Model performance was assessed independently within each space using the Pearson correlation and mean squared error (MSE) between the corresponding empirical and simulated connectivity matrices. Both measures remained stable across treatment groups and sessions (**Supplementary Tables S1 and S2**). The resulting subject- and session-specific GEC and GECVO matrices, together with empirical vortex-space dynamics, were used in the subsequent analyses summarised in **Figure 1D**.

### Differential sensitivity to treatment-related brain reconfiguration

We next used supervised machine learning to compare the sensitivity of vortex and signal spaces across four classification analyses (**Figure 2A**). Subject- and session-specific GECVO and GEC matrices were classified separately to distinguish pre- from post-treatment brain states within each of the psilocybin and escitalopram groups, discriminate between the post-treatment states associated with the two interventions, and predict subsequent clinical response to escitalopram from baseline connectivity. The response analysis was restricted to the escitalopram group, which had a more balanced distribution of responders and non-responders.

**Figure 2.**
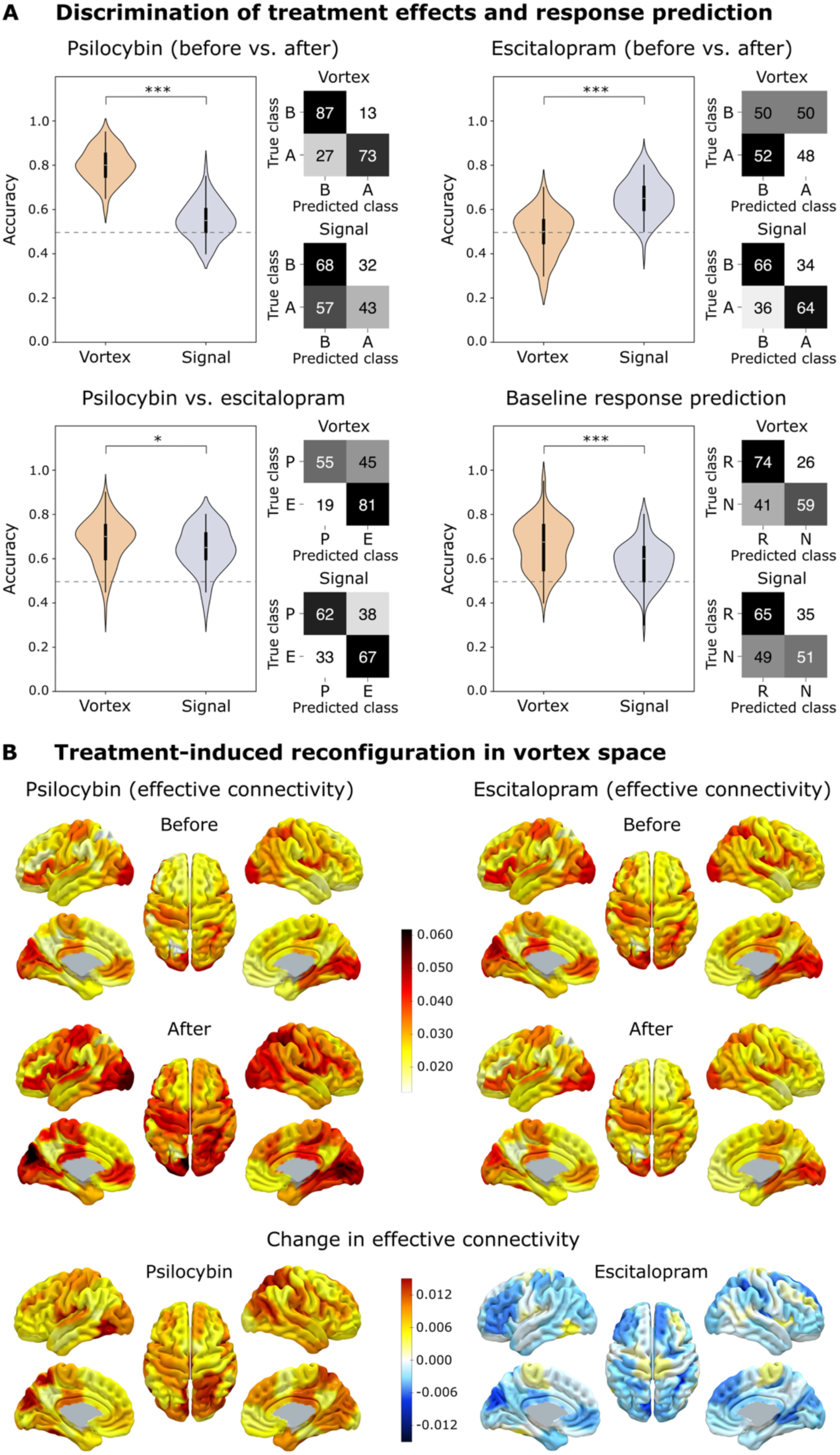
Distinct patterns of brain reconfiguration following psilocybin and escitalopram treatment and prediction of clinical response. **A.** Machine-learning classifiers were trained independently using subject-specific vortex-space (GECVO) and signal-space (GEC) effective-connectivity matrices to distinguish pre- from post-treatment brain states within each intervention, discriminate between the interventions at post-treatment, and predict subsequent response to escitalopram from baseline connectivity. Violin plots show distributions of cross-validated accuracy, dashed lines indicate chance performance, and confusion matrices show class-specific classification rates. Vortex-space models achieved higher accuracy for psilocybin pre–post classification, treatment discrimination and baseline response prediction, whereas signal-space models performed better for escitalopram pre–post classification. Asterisks indicate significant differences between representations (*P < 0.05, **P < 0.01, and ***P < 0.001). **B.** Group-average GECVO is mapped onto the Schaefer-100 parcellation in MNI152 space before and after each intervention, with the bottom row showing post-minus-pre changes. Brain renderings show widespread positive changes following psilocybin, compared with smaller, spatially heterogeneous and predominantly negative changes following escitalopram.

For each classification, connections were ranked by between-class discriminability using Wilcoxon tests with 10,000 permutations. Paired tests were used for longitudinal comparisons and unpaired tests for treatment and response classification. Progressively larger subsets of the highest-ranked connections were evaluated using the cross-validation procedure described in *Methods*. Both feature ranking and subset selection were performed independently within each training fold to prevent information leakage. Cross-validated accuracy summarised overall performance, while confusion matrices showed class-specific classification rates. For each classification, accuracy was compared between vortex and signal spaces using Wilcoxon tests (10,000 permutations); all *P* values reported for these comparisons derive from those tests.

The space yielding higher accuracy depended on the classification. Psilocybin pre–post classification reached an accuracy of 0.80 ± 0.25 (mean ± s.d.) in vortex space and 0.55 ± 0.25 in signal space, using 2,000 and 100 connections, respectively (*P* < 0.001). The corresponding confusion matrices showed clearer separation between pre- and post-treatment states in vortex space. By contrast, escitalopram pre–post classification reached an accuracy of 0.65 ± 0.28 in signal space, with balanced classification of both states, compared with 0.49 ± 0.29 in vortex space, in both cases using 10 connections (*P* < 0.001).

At post-treatment, classification of psilocybin versus escitalopram participants was slightly more accurate in vortex space. Accuracy reached 0.68 ± 0.31 in vortex space and 0.64 ± 0.32 in signal space, with both models using 2,000 connections (*P* = 0.017). This difference was largely driven by more accurate classification of escitalopram-treated participants. Similarly, subsequent response to escitalopram was classified from baseline connectivity with an accuracy of 0.66 ± 0.34 in vortex space and 0.58 ± 0.31 in signal space, using 20 and 750 connections, respectively (*P* < 0.001). The confusion matrices showed higher class-specific accuracy for responders than for non-responders.

To characterise the spatial distribution of treatment-related changes, regional global brain connectivity (GBC) was calculated as each region’s mean effective connectivity with all others. Within-treatment longitudinal effects were assessed using paired Wilcoxon tests with 10,000 permutations, whereas post-minus-pre changes were compared between treatments using unpaired tests. *P* values were corrected across the 100 regions using the Benjamini–Hochberg false discovery rate (FDR) procedure. In vortex space, psilocybin produced significant increases from pre- to post-treatment in 76 of 100 regions, spanning all seven canonical Yeo networks (Yeo et al., 2011), whereas no changes survived FDR correction following escitalopram. Post-minus-pre changes differed significantly between treatments in 64 of 100 regions (**Supplementary Table S3**). In signal space, conversely, no regional effects survived correction for either longitudinal comparison or the between-treatment contrast (**Supplementary Table S4**).

The spatial distribution of these effects in vortex space is shown in **Figure 2B**. Psilocybin produced widespread and consistently positive changes in effective connectivity. The largest changes were observed in dorsal-attention, somatomotor and visual regions, with additional involvement of frontoparietal control (hereafter, control) and salience/ventral-attention regions (**Supplementary Table S5**). Escitalopram, by contrast, showed smaller and more spatially heterogeneous changes that were predominantly negative, distributed mainly across dorsal-attention, control, default-mode and visual regions, although none survived regional FDR correction (**Supplementary Table S6**). The largest between-treatment differences similarly extended across multiple networks, particularly dorsal-attention, visual, salience/ventral-attention and control systems (**Supplementary Table S7**).

### Baseline effective connectivity associated with subsequent clinical improvement

We next examined whether baseline effective-connectivity patterns were associated with subsequent clinical improvement following psilocybin and escitalopram treatment (**Figure 3**). Separately for each treatment and space, regional baseline GBC was correlated with clinical improvement, defined as the reduction in BDI score. Across a range of correlation thresholds, regions showing positive or negative associations were selected and combined into two participant-specific connectivity scores, which were entered into a linear model adapted from connectome-based predictive modelling (Shen et al., 2017). Model fit was quantified in-sample as the correlation between observed and fitted improvement, with higher values indicating that the selected baseline connectivity pattern explained more interindividual variation in symptom reduction. For each space, we retained the threshold yielding the strongest in-sample correlation.

**Figure 3.**
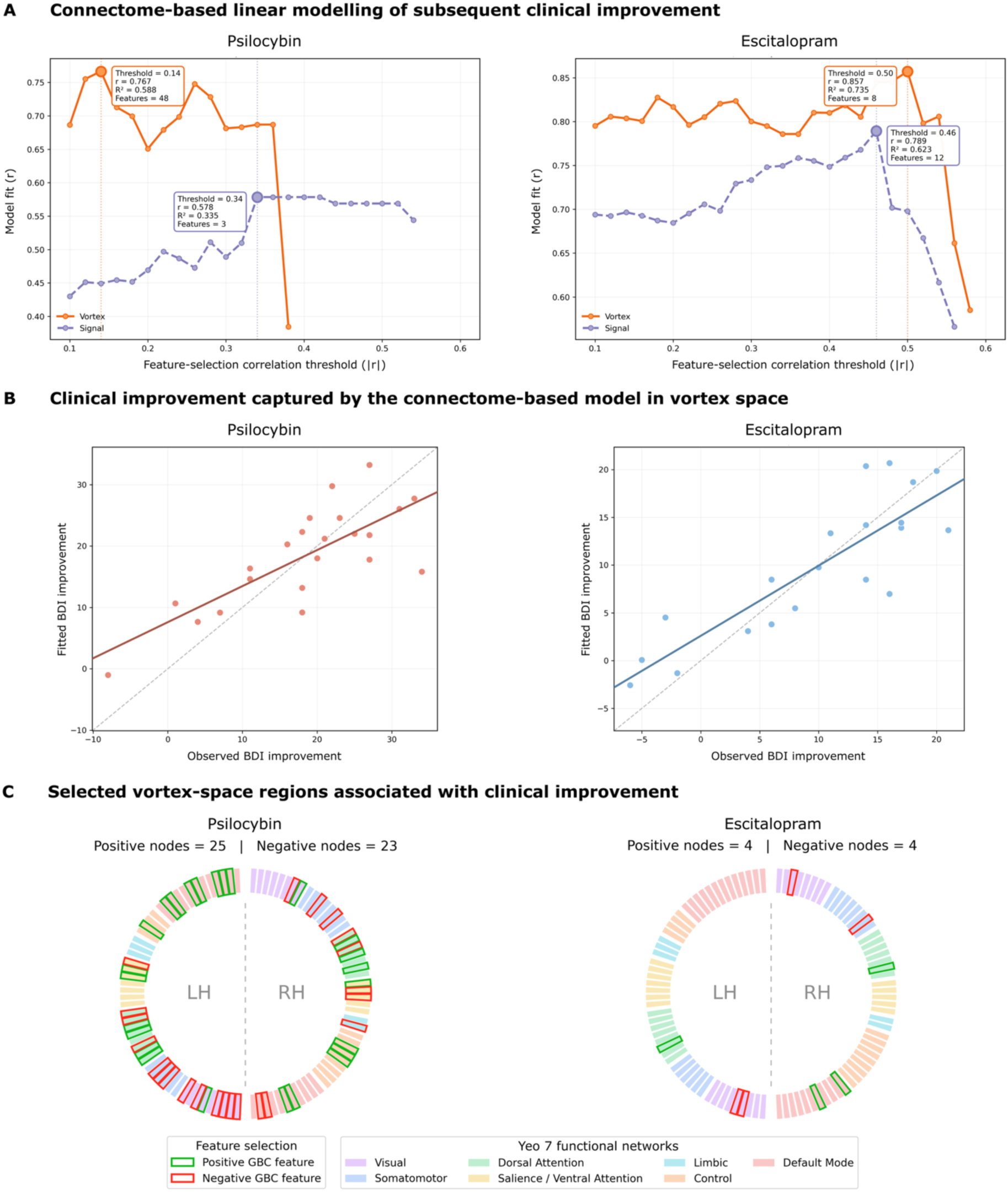
Baseline effective connectivity associated with subsequent clinical improvement. Clinical improvement was defined as pre-treatment minus post-treatment BDI score, such that higher values indicate greater symptom reductions. **A.** Connectome-based linear models relating baseline regional GBC to subsequent clinical improvement following psilocybin or escitalopram treatment. In-sample model fit is shown across feature-selection correlation thresholds (|r|) for vortex and signal spaces. The optimal threshold was defined as that yielding the highest correlation between observed and fitted improvement. Vortex-space models achieved higher maximum fits than signal-space models for both interventions. **B.** Observed improvement is plotted against the values fitted by the optimal vortex-space model for each intervention. Solid lines show linear fits, and the dashed diagonal indicates perfect agreement between observed and fitted values. **C.** Regional GBC features selected by the optimal vortex-space models. Green and red outlines indicate features positively and negatively associated with clinical improvement, respectively, while regional colours indicate Yeo 7-network assignment of each Schaefer-100 parcel. Regions are arranged by hemisphere; LH, left hemisphere; RH, right hemisphere.

The strongest in-sample associations for both treatments were obtained in vortex space (**Figure 3A**). For psilocybin, the optimal vortex-space model reached *r* = 0.77 (*R*² = 0.59), using 48 regional GBC features selected at |*r*| = 0.14. The corresponding signal-space model reached *r* = 0.58 (*R*² = 0.34), using three features selected at |*r*| = 0.34. A similar pattern was observed for escitalopram: the optimal vortex-space model reached *r* = 0.86 (*R*² = 0.74), using eight features selected at |*r*| = 0.50, whereas the signal-space model reached *r* = 0.79 (*R*² = 0.62), using 12 features selected at |*r*| = 0.46. The observed-versus-fitted relationships for the optimal vortex-space models are shown in **Figure 3B**.

At the optimal thresholds, the selected regional GBC features revealed distinct spatial patterns across treatments (**Figure 3C**). For psilocybin, 48 regions were selected, comprising 25 positive and 23 negative associations distributed bilaterally across all seven canonical Yeo networks (Yeo et al., 2011). By contrast, the escitalopram model selected only eight regions, equally divided between positive and negative associations and restricted to the visual, somatomotor, dorsal-attention, control and default-mode networks.

### Treatment-related reconfiguration of network modularity

Global network reconfiguration can be quantified through modularity, which captures the extent to which the brain is organised into internally cohesive and externally segregated communities (Newman & Girvan, 2004). Accordingly, we identified community structure from subject- and session-specific GECVO and GEC matrices and quantified modularity in vortex and signal spaces before and after psilocybin and escitalopram treatment (**Figure 4A**). Longitudinal changes were assessed within each intervention, and treatment-related changes, defined as post-minus-pre modularity, were compared between psilocybin and escitalopram.

**Figure 4.**
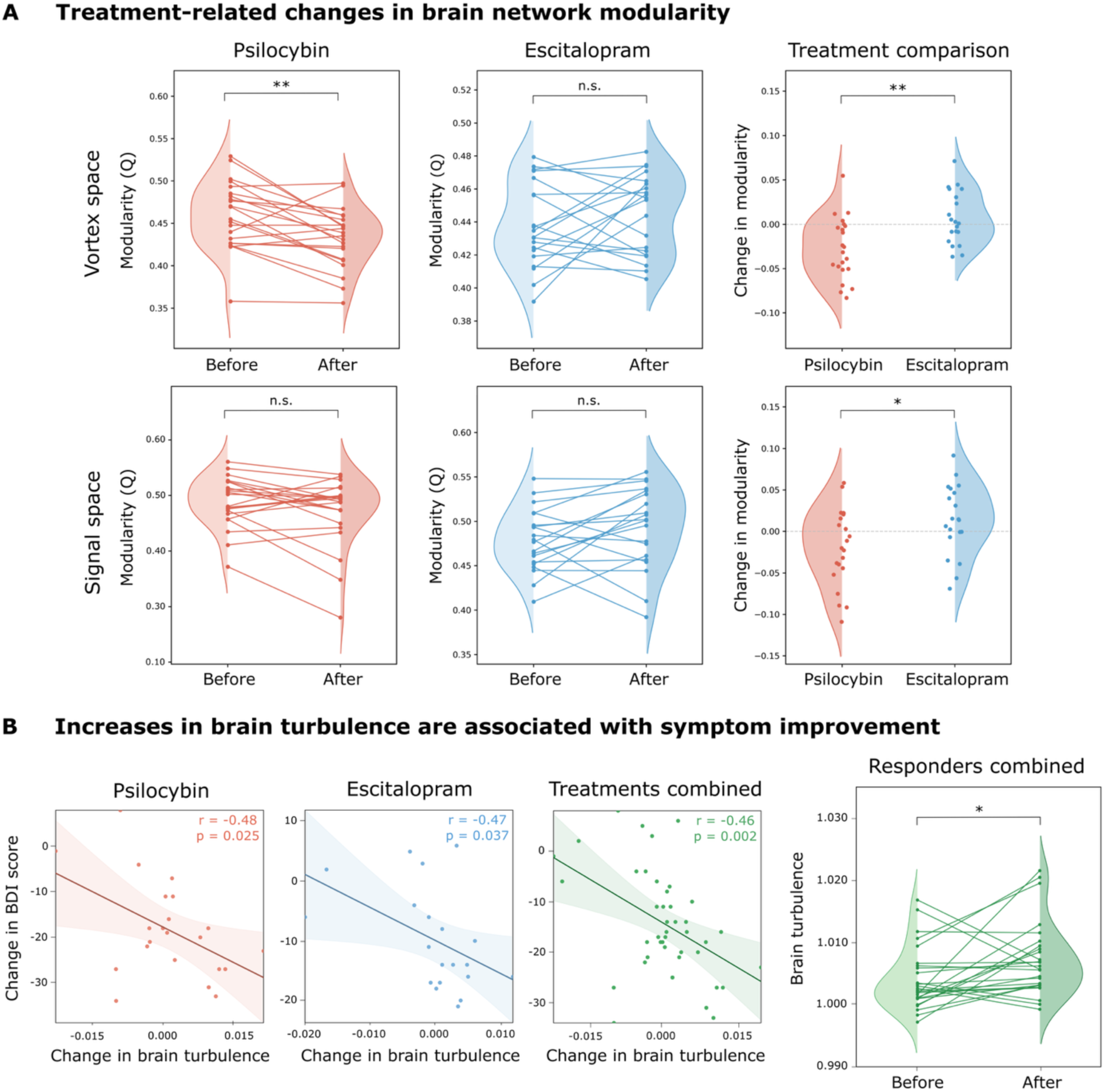
Treatment-related network reconfiguration and brain turbulence associated with clinical improvement. **A.** Network modularity is shown before and after psilocybin and escitalopram treatment in vortex and signal spaces, together with post-minus-pre changes compared between treatments. Lines connect repeated measures. Psilocybin reduced modularity, significantly in vortex space, whereas escitalopram showed no significant change. Between-treatment differences were significant in both representations and clearer in vortex space. **B.** Pearson correlations between post-minus-pre changes in brain turbulence and BDI score are shown for each treatment and for the pooled sample. Negative BDI changes indicate reductions in depressive symptoms. Larger increases in turbulence were associated with greater clinical improvement, and pooled responders (n = 26) showed a significant post-treatment increase in turbulence. Asterisks indicate statistical significance (*P < 0.05, **P < 0.01, and ***P < 0.001); n.s., not significant.

Psilocybin was associated with a significant reduction in network modularity in vortex space (*P* = 0.002, *d* = −0.79), but not in signal space (*P* = 0.133, *d* = −0.40). Escitalopram showed no significant longitudinal change in either vortex (*P* = 0.442, *d* = 0.22) or signal space (*P* = 0.125, *d* = 0.34). Nevertheless, the difference in modularity change (i.e., post-minus-pre) between the two treatments was significant in both spaces, with a larger effect in vortex space (*P* = 0.003, |*d*| = 1.05) than in signal space (*P* = 0.042, |*d*| = 0.74).

### Brain turbulence associated with clinical improvement

Brain turbulence captures the spatiotemporal variability of local neural synchronisation, and has been proposed as an index of the brain’s dynamical flexibility and capacity for reconfiguration (Deco et al., 2025). For each participant and session, empirical amplitude turbulence was quantified directly from the variability of pairwise co-fluctuations in local synchronisation across regions and time (*Methods*). We then examined whether post-minus-pre changes in turbulence were associated with changes in BDI score following psilocybin and escitalopram treatment (**Figure 4B**), with more negative BDI changes indicating greater symptom reduction.

Increases in brain turbulence were significantly correlated with greater reductions in depressive symptoms following psilocybin (*r* = −0.48, *P* = 0.025) and escitalopram (*r* = −0.47, *P* = 0.037). This relationship remained significant when both treatment groups were combined (*r* = −0.46, *P* = 0.002). To better understand the changes in turbulence associated with clinical response, we examined longitudinal changes among participants meeting the prespecified response criterion. Notably, responders pooled across both interventions (*n* = 26) showed a significant increase in brain turbulence after treatment (*P* = 0.018, *d* = 0.46).

## Discussion

In this study, we investigated how psilocybin and escitalopram reshape large-scale brain dynamics in patients with MDD using signal- and vortex-space whole-brain models. By capturing the turbulent spatiotemporal organisation of brain activity, vortex space was more sensitive than signal space to treatment-related brain reorganisation in most analyses. Importantly, psilocybin and escitalopram had distinct effects on brain dynamics despite their shared therapeutic aim. These differences extended from effective connectivity to global network organisation. Furthermore, baseline connectivity carried information about subsequent clinical improvement, while changes in empirical turbulence were associated with symptom reduction in both groups. Overall, our findings extend previous work investigating the distinct patterns of brain reconfiguration following psilocybin and escitalopram (Dagnino et al., 2026; Deco et al., 2024; Socoró-Garrigosa et al., 2025) and bring vortex-space whole-brain modelling (Deco et al., 2026) into computational psychiatry. Although replication in larger cohorts and across other interventions is necessary to establish generalisability, this framework offers a promising route towards using neuroimaging to better understand the dynamical mechanisms underlying neuropsychiatric disorders and their successful treatment.

### Vortex-space modelling shows greater sensitivity to treatment-related brain reconfiguration

Signal and vortex spaces capture distinct features of brain dynamics. Conventional signal-space modelling characterises interactions between mean regional BOLD signals, whereas vortex-space modelling characterises interactions between mesoscopic patterns of local synchronisation. Their relative sensitivity therefore depended on the information examined rather than one space being uniformly superior. The clearest advantage of vortex space concerned psilocybin: vortex-space effective connectivity classified pre- and post-treatment states more accurately (**Figure 2A**), while widespread increases in regional GBC across all seven canonical networks (**Supplementary Tables S3 and S4**) and a significant reduction in modularity (**Figure 4A**) were detected only in this space. Together, these findings suggest that changes following psilocybin were strongly expressed in the distributed interactions between local synchronisation patterns, a level of organisation not directly represented in signal space. This interpretation is consistent with evidence that acute psilocybin increases the spatiotemporal variability of local synchronisation in healthy individuals, particularly over longer spatial scales (Cruzat et al., 2022).

The same sensitivity extended to the direct comparison of psilocybin and escitalopram. Vortex space more accurately distinguished their post-treatment states (**Figure 2A**), detected differences between the treatments in post-minus-pre regional GBC that were not observed in signal space (**Supplementary Tables S3 and S4**), and showed a larger effect size for the between-treatment difference in modularity change (**Figure 4A**). Thus, the distinct signatures associated with psilocybin and escitalopram appear to be more clearly expressed in the organisation of local synchronisation patterns. Previous work supports the relevance of this level of organisation for characterising different brain states, showing that turbulence-based measures distinguish resting wakefulness, meditation, deep sleep and post-coma states of consciousness (Escrichs et al., 2022) and that modelling vortex interactions discriminates rest from cognitive tasks and subtle task components (Deco et al., 2026).

Notably, vortex space also captured clinically relevant interindividual variability. Baseline GECVO classified subsequent response to escitalopram more accurately than baseline GEC (**Figure 2A**), while connectome-based models using baseline vortex-space GBC showed stronger in-sample associations with clinical improvement for both treatments (**Figure 3A**). These findings are consistent with evidence that baseline turbulence predicts subsequent response to escitalopram or duloxetine in MDD (Escrichs et al., 2025). Although based on different measures—effective interactions between local synchronisation patterns here, and their variability in the previous study—both studies suggest that the organisation of local synchronisation before treatment carries information relevant to individual differences in antidepressant response.

Signal space nevertheless classified pre- and post-treatment states following escitalopram more accurately (**Figure 2A**). This indicates that longitudinal effects following escitalopram were more discriminable in effective connectivity derived from regional BOLD activity, whereas response-related differences were more evident in the organisation of local synchronisation. It is tempting to speculate that this dissociation reflects the substantial interindividual variability in escitalopram response—with vortex space preferentially capturing response-related heterogeneity and signal space capturing a broader longitudinal signature of escitalopram treatment. However, further work will be required to test this hypothesis directly. The observed dissociation therefore reinforces the conclusion that signal and vortex spaces capture complementary aspects of treatment-related brain dynamics.

### Distinct dynamical reconfiguration following psilocybin and escitalopram

Regional analyses in vortex space revealed markedly distinct patterns of reconfiguration following psilocybin and escitalopram. Psilocybin produced widespread increases in regional GBC spanning all seven canonical Yeo networks (Yeo et al., 2011). Conversely, escitalopram showed smaller, spatially heterogeneous and predominantly negative changes, although none survived FDR correction. Consistent with this contrast, changes differed between treatments in 64 of 100 regions (**Supplementary Table S3**). This difference in spatial extent parallels previous whole-brain modelling of the same trial, which revealed widespread regional hierarchical reconfiguration following psilocybin but comparatively limited changes following escitalopram (Deco et al., 2024).

The largest psilocybin-related increases involved dorsal-attention, somatomotor and visual regions, with additional contributions from control and salience/ventral-attention systems (**Supplementary Table S5**). This distribution is consistent with evidence that acute psychedelic administration strengthens connectivity between visual, somatomotor and dorsal-attention systems and transmodal default-mode and frontoparietal networks (Girn et al., 2026). For escitalopram, the largest changes involved dorsal-attention, control, default-mode and visual regions (**Supplementary Table S6**), aligning with previous evidence that escitalopram modifies effective connectivity involving the dorsal-attention, default-mode and salience networks (Wang et al., 2024). The strongest between-treatment differences were distributed primarily across dorsal-attention, visual, salience/ventral-attention and control systems (**Supplementary Table S7**). Thus, the distinction between the treatments may lie less in which functional systems are involved and more in the magnitude, spatial extent and direction of their reconfiguration.

At the global network level, psilocybin was associated with a significant reduction in vortex-space modularity, whereas no significant longitudinal change was observed following escitalopram. The two treatments also differed significantly in their post-minus-pre change in modularity (**Figure 4A**). The reduction in modularity following psilocybin indicates a shift towards a less segregated network organisation. This pattern aligns with Daws et al. (2022), who reported reduced FC modularity after psilocybin treatment but no comparable change with escitalopram. Recent whole-brain modelling incorporating temporal asymmetry likewise found decreased GEC modularity after psilocybin, but reported an increase following escitalopram (Dagnino et al., 2026). This discrepancy may reflect differences in the study samples, brain parcellation, or the modelling framework. More broadly, the consistent reduction in modularity after psilocybin treatment parallels findings during acute LSD administration (Tagliazucchi et al., 2016), perhaps reflecting a therapeutically relevant carry-over of acute psychedelic effects.

These results form part of a broader pattern emerging across complementary whole-brain modelling analyses of the trial analysed here. Deco et al. (2024) found that psilocybin reduced the global directedness of the functional hierarchy, whereas escitalopram increased it. Using in silico perturbations, Socoró-Garrigosa et al. (2025) showed that psilocybin increased the brain’s average susceptibility to perturbation, whereas escitalopram reduced it. Dagnino et al. (2026) similarly found opposing changes in non-equilibrium dynamics, with increased deviation from the fluctuation-dissipation theorem following psilocybin and decreased deviation following escitalopram. Although these approaches quantify distinct aspects of brain dynamics, they consistently reveal markedly different effects of psilocybin and escitalopram. Overall, this convergence suggests that similar therapeutic outcomes may arise from distinct and, in some respects, opposing forms of large-scale brain reorganisation.

### Clinical relevance of effective connectivity and brain turbulence

Importantly, baseline GECVO carried clinically relevant information about subsequent treatment outcome. In the escitalopram group, where the balance of responders and non-responders allowed cross-validated classification, subsequent clinical response could be predicted from baseline connectivity significantly above chance (**Figure 2A**). Beyond this binary classification, baseline regional GBC showed strong in-sample associations with subsequent clinical improvement for both interventions (**Figure 3**). These results align with earlier studies linking pre-treatment brain connectivity to subsequent clinical outcomes, including response to escitalopram (DeMaster et al., 2022) and remission following antidepressant treatment (Korgaonkar et al., 2019). Whole-brain modelling has likewise highlighted the clinical relevance of pre-treatment brain dynamics. In the same trial, baseline hierarchical organisation predicted subsequent response to escitalopram (Deco et al., 2024), while a separate study of psilocybin identified pre-treatment dynamical configurations associated with later psilocybin response (Vohryzek et al., 2024). Our results extend this evidence to vortex-space effective connectivity. Together, these findings suggest that baseline connectivity may help explain individual differences in subsequent antidepressant outcomes. Further work is needed to establish the robustness and generalisability of these associations and evaluate their potential to inform personalised treatment.

The regional configurations associated with clinical improvement also differed across treatments (**Figure 3C**). For psilocybin, the selected baseline regions were distributed bilaterally across all seven canonical Yeo networks (Yeo et al., 2011), whereas the escitalopram model showed a more spatially restricted pattern involving visual, somatomotor, dorsal-attention, control and default-mode systems. These results support the view that clinical improvement following the two interventions may be associated with distinct baseline configurations of large-scale brain organisation. However, this spatial contrast should be interpreted descriptively, as each model was evaluated in-sample at its own optimal feature-selection threshold.

Treatment-related changes in empirical brain turbulence were also clinically relevant. Greater increases in turbulence were associated with larger reductions in depressive symptoms following both psilocybin and escitalopram, and this association remained significant when all participants were considered together (**Figure 4B**). Consistently, responders pooled across interventions exhibited a significant post-treatment increase in turbulence. This convergence across treatments contrasts with the divergent patterns of brain reconfiguration observed in the preceding analyses, suggesting that increased brain turbulence may represent a shared correlate of clinical improvement. Future work should determine whether increased turbulence reflects a common underlying mechanism or arises through treatment-specific processes, and whether it contributes to symptom reduction, emerges as a consequence of improvement, or accompanies it without being mechanistically involved. Replication in independent cohorts and across interventions will be necessary to establish whether changes in brain turbulence could serve as a general and longitudinal marker of mental health improvement.

### Potential limitations and future directions

Our study has several limitations. The whole-brain models were expressed over the 100 cortical regions of the Schaefer-100 parcellation, which provided a common and computationally tractable scale for comparing signal and vortex spaces. The findings are therefore constrained to this resolution and do not capture subcortical regions that may be relevant to depression treatment. The models also used a group-average structural connectome from an independent healthy cohort because individual structural connectivity was unavailable. Nevertheless, this template provided a common anatomical starting point, with GEC and GECVO optimisation adapting the weights of its existing connections to the empirical data from each participant and session. Lastly, the relatively small sample limits statistical power and the stability of individual-level clinical analyses. The imbalance between responders and non-responders in the psilocybin group restricted cross-validated response classification to escitalopram, while the connectome-based models were evaluated in-sample and should therefore be regarded as associative rather than predictive. Consequently, some of the findings may not be representative of the broader population of individuals with MDD treated with psilocybin or escitalopram. Replication in larger independent cohorts would increase statistical power and improve the robustness and generalisability of the conclusions.

### Conclusion

Overall, modelling vortex interactions provides a sensitive description of how interventions reshape brain dynamics in depression. This arises from combining the generative principles of whole-brain modelling with an explicit representation of the brain’s turbulent dynamics, thereby capturing a mesoscopic level of organisation not directly represented in regional BOLD signals. Across most analyses, vortex space was more sensitive to treatment-related changes than conventional signal space. Directly addressing our main research question, psilocybin and escitalopram appear to alleviate depressive symptoms through predominantly distinct patterns of brain reconfiguration. Psilocybin was associated with widespread increases in GECVO and reduced network modularity, whereas no comparable changes were observed following escitalopram. Importantly, baseline patterns of GECVO carried information about subsequent treatment outcome and predicted response to escitalopram significantly above chance. Changes in brain turbulence were also associated with clinical improvement across both treatments, raising the possibility that turbulence could provide a useful general and longitudinal marker of symptom reduction. Larger independent studies are now needed to establish the generalisability and predictive value of this framework, together with its potential contribution to more personalised treatment strategies.

## Methods

### Clinical trial and participants

Data were obtained from a phase II double-blind randomised controlled trial comparing psilocybin and escitalopram in patients with MDD (ClinicalTrials.gov ID: NCT03429075), as previously reported (Carhart-Harris et al., 2021; Daws et al., 2022). A schematic overview of the trial design is shown in **Figure 1A**. Only the aspects relevant to the present study are illustrated; other conditions and measures from the original trial are not included.

The study was conducted at the Imperial Clinical Research Facility (London, UK) under sponsorship from Imperial College London, with approvals from the NHS Research Ethics Committee (17/LO/0389), the Health Research Authority, and the Medicines and Healthcare Products Regulatory Agency. The protocol was implemented under a UK Home Office Schedule 1 licence. All participants provided written informed consent in accordance with institutional and national ethical standards.

Eligible patients had a confirmed diagnosis of unipolar MDD and scored ≥16 on the 21-item Hamilton Depression Rating Scale. Exclusion criteria included psychotic or bipolar disorders, serious medical illness, MRI or SSRI contraindications, a positive pregnancy test, or a history of severe suicide attempts. Treatment resistance was not used as an inclusion or exclusion criterion. All participants completed structured screening, clinical interviews, and comprehensive psychiatric and medical assessments.

A total of 59 patients were randomly assigned to either psilocybin (n = 30) or escitalopram (n = 29). After quality control and motion exclusion, 22 participants from the psilocybin group (mean age = 41.9 ± 11.0 years; 14 men, 8 women) and 20 from the escitalopram group (mean age = 38.7 ± 11.0 years; 14 men, 6 women) were retained for the neuroimaging analyses. Resting-state fMRI data were available before and after treatment. Prior psychedelic exposure was documented in 31% of participants in the psilocybin arm and 24% in the escitalopram arm. Demographic information is summarised in **Supplementary Table S8**.

### Treatment protocol

The trial comprised two dosing sessions and daily oral administration over six weeks and one day. On the first dosing day (DD1), participants received either 25 mg of psilocybin (psilocybin arm) or a 1 mg dose of psilocybin, considered pharmacologically negligible (escitalopram arm). All participants were informed that they would receive psilocybin but were blinded to the administered dose. A second identical dosing session (DD2) occurred three weeks later. From the day following DD1, participants took one capsule daily for the first three weeks and two capsules daily thereafter. Capsules contained either inert placebo in the psilocybin arm or escitalopram (10 mg for the first three weeks and 20 mg thereafter) in the escitalopram arm. Participants remained in their assigned treatment arm throughout the study. Further methodological and clinical details are available in the primary publication of the trial (Carhart-Harris et al., 2021).

### Neuroimaging data

Resting-state fMRI data were acquired before and after treatment using a 3T Siemens Tim Trio scanner with a 32-channel head coil (Invicro, London, UK). Each session lasted approximately ten minutes and consisted of 480 T2*-weighted echo-planar volumes (TR = 1,250 ms; TE = 30 ms; voxel size = 3 mm isotropic; 44 axial slices; flip angle = 70°; GRAPPA acceleration = 2; bandwidth = 2,232 Hz per pixel). Participants were instructed to keep their eyes closed during acquisition.

Preprocessing was performed using a standard in-house pipeline based on FSL, AFNI, FreeSurfer, and ANTs, following established procedures (Avants et al., 2009; Cox, 1996; Dale et al., 1999; Smith et al., 2004). Steps included de-spiking, slice-timing and motion correction, skull stripping, rigid-body registration to individual anatomy, nonlinear normalisation to Montreal Neurological Institute (MNI) space, scrubbing, band-pass filtering, and regression of motion, tissue, venous, and white-matter confounds. Motion artefacts were carefully inspected, and no systematic between-group differences in motion were detected, as reported previously (Daws et al., 2022). BOLD signals were parcellated using the Schaefer-1000 atlas, which divides the cortex into 1,000 regions of interest (500 per hemisphere) based on intrinsic functional connectivity patterns (Schaefer et al., 2018). These time series served as the high-resolution basis for the subsequent signal- and vortex-space representations, which were ultimately analysed at the Schaefer-100 level.

To define the anatomical scaffold for the whole-brain models, we used a group-average structural connectivity (SC) matrix derived from high-quality multi-shell diffusion MRI data from 32 healthy participants from the Human Connectome Project (HCP). Participants were scanned at Massachusetts General Hospital (MGH) on a customised 3-T Siemens Connectom scanner, a modified 3-T Skyra system, using a 64-channel head coil. For each participant, a T1-weighted MPRAGE image was acquired at 1.0-mm isotropic resolution and a high-resolution T2-weighted SPACE image at 0.7-mm isotropic resolution, together with multi-shell diffusion data acquired over approximately 89 minutes (Setsompop et al., 2013).

Diffusion data were reconstructed using generalised q-sampling imaging in DSI Studio, with 200,000 streamlines per participant (Horn et al., 2017). White-matter masks were derived from segmentations of the T2-weighted images co-registered to the diffusion b₀ image using SPM12. Streamlines were normalised to MNI space with Lead-DBS (Horn & Blankenburg, 2016) and parcellated according to the Schaefer-100 atlas (Schaefer et al., 2018). To minimise false-positive connections, we employed tractography algorithms validated in previous benchmarking work, achieving a 92% valid-connection rate across independent submissions (Maier-Hein et al., 2017). The final SC matrix was symmetrised and normalised, with its diagonal set to zero to remove self-connections.

### Clinical data

Depressive symptom severity was assessed using the Beck Depression Inventory (BDI) (Beck et al., 1961). This self-rated questionnaire captures a broad spectrum of depressive symptoms, with higher scores indicating greater symptom severity. Assessments were obtained at baseline and after 2, 4, and 6 weeks of treatment. Although the Quick Inventory of Depressive Symptomatology (QIDS) was the prespecified primary outcome of the original clinical trial (Carhart-Harris et al., 2021), the present study used the BDI to quantify symptom change and define clinical response to maintain consistency with previous neuroimaging and whole-brain modelling analyses of the same dataset (Daws et al., 2022; Deco et al., 2024).

Clinical response was defined as a ≥ 50% reduction in BDI score from baseline to week 6, and remission as a week-6 BDI score ≤ 9. Among participants retained for the neuroimaging analyses, 18 of 22 participants in the psilocybin group and 8 of 20 in the escitalopram group met the response criterion, compared with 4 and 12 non-responders, respectively. BDI remission rates were 64% in the psilocybin group and 30% in the escitalopram group. Response classifications were used in subsequent analyses.

### Vortex space representation

An intuitive approach to characterising brain dynamics is to assess the degree of global phase synchrony across the cortex (Hancock et al., 2022; Sanz Perl, Zamora-Lopez, et al., 2023). This can be quantified using the global Kuramoto order parameter, defined as:

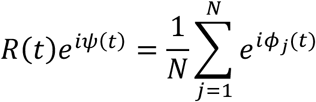

where *N* denotes the number of brain regions, and *φ*_(_(*t*) is the instantaneous phase of the BOLD signal at region *j*. The magnitude *R*(*t*) ∈ [0,1] reflects the level of global synchronisation at each time point, with higher values indicating stronger coherence across the brain. However, in resting-state fMRI data, *R*(*t*) typically remains below unity, revealing that the brain is not fully synchronised. Instead, it exhibits substantial local fluctuations in phase alignment, with patterns of partial synchrony that evolve across space and time (Deco et al., 2017).

To capture these localised patterns of coordination, brain dynamics must be represented at an intermediate, mesoscopic scale. This is accomplished through the construction of vortex space, a representation that captures the local synchrony of functionally coherent groups of brain regions (Deco et al., 2026).

### Processing and phase extraction of BOLD signals

BOLD time series were extracted from 1,000 cortical parcels defined by the Schaefer-1000 atlas, which provides a fine-grained functional parcellation of the cortex (Schaefer et al., 2018). Each regional signal was linearly detrended to remove slow drifts and z-scored to normalise across regions. A second-order zero-phase Butterworth band-pass filter (0.008–0.08 Hz) was then applied to retain slow oscillatory components characteristic of resting-state activity, while attenuating low-frequency drifts and high-frequency physiological noise (Fox et al., 2005; Lowe et al., 1998). To minimise edge artefacts, the first and last ten volumes of the filtered signal were discarded.

Then, the instantaneous phase of each regional BOLD signal was computed using the Hilbert transform. The analytic signal for each region *j* was defined as:

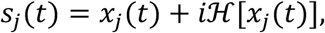

where *x*_(_(*t*) denotes the filtered BOLD time series and ℋ[*x*_(_(*t*)] its Hilbert transform (Sanz Perl, Zamora-Lopez, et al., 2023). The phase time series *φ*_(_(*t*) = arg(*s*_(_(*t*)) was then obtained and used as input for computing local synchrony in the empirical vortex space.

### Kuramoto vorticity and local synchrony

Turbulence arises from fluctuations in local phase synchrony that propagate across space and time. This phenomenon, first described in coupled oscillator systems (Kawamura et al., 2007), and later extended to brain dynamics (Deco et al., 2025, 2026; Deco & Kringelbach, 2020), supports efficient information transfer across scales (Deco et al., 2023). Within the Kuramoto framework (Kuramoto, 1984), these interactions are formalised by the local Kuramoto order parameter, which represents a spatial average of the complex phase factors of coupled oscillators within a neighbourhood.

To characterise these interactions at the mesoscale, cortical activity was coarse-grained into 100 macroregions following the Schaefer-100 functional atlas, each comprising spatially contiguous parcels from the Schaefer-1000 parcellation (Schaefer et al., 2018). Within each macroregion σ, local synchrony was quantified using the local Kuramoto order parameter:

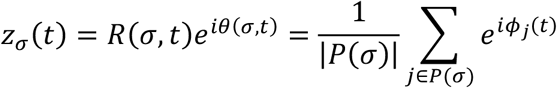

where *φ*_(_(*t*) denotes the instantaneous phase of parcel *j* at time *t*. The complex Kuramoto order parameter *z_σ_*(*t*) ∈ ℂ captures the degree of phase alignment among parcels within macroregion σ. Its modulus *R*(*σ*, *t*) = |*z_σ_*(*t*)|, also referred to as Kuramoto vorticity, quantifies the strength of local synchrony, ranging from 0 (complete desynchronisation within the region) to 1 (perfect phase locking), while its argument *θ*(*σ*, *t*) = arg(*z_σ_*(*t*)) represents the dominant local phase.

Together, *R*(*σ*, *t*) and *θ*(*σ*, *t*) describe the amplitude and phase components of local coordination, providing a natural mesoscale representation of brain activity. In this formulation, each macroregion can be interpreted as a dynamic vortex, characterised by an evolving amplitude and phase that fluctuate over time and interact across space. This defines the vortex space representation of brain dynamics, which serves as the empirical foundation for subsequent whole-brain modelling.

### Brain turbulence

To quantify the spatiotemporal variability of local synchrony, we computed a measure of brain turbulence from the local synchrony field. For each macroregion, Kuramoto vorticity, *R*(*σ*, *t*), was first normalised across time to remove baseline offsets. Pairwise co-fluctuations were then computed at each time point as the element-wise product of the normalised amplitudes of each region pair, *R*^∼^(*σ*, *t*)*R*^∼^(*σ*^1^, *t*), yielding a time-resolved matrix of synchrony co-fluctuations.

Kuramoto turbulence, hereafter referred to as amplitude turbulence, was defined as the standard deviation of these co-fluctuation values across all region pairs and time points:

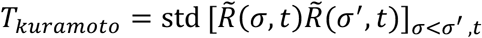

Higher values therefore indicate greater spatiotemporal variability in local synchrony across the cortex.

### Vortex-space functional connectivity

To characterise the coordination between distinct vortices, we computed the vortex-space functional connectivity (FV) matrix. For each pair of macroregions (*σ*, *σ*^1^), this matrix quantifies the strength of their co-fluctuations in local synchrony while maintaining phase alignment. Specifically, it was defined as:

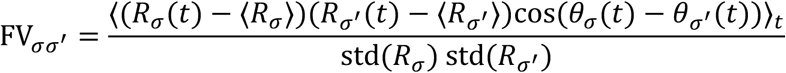

where *R_σ_*(*t*) and *θ_σ_*(*t*) denote the vorticity amplitude and phase of macroregion *σ* at time *t*, respectively. This formulation extends conventional correlation measures by integrating both amplitude variability and phase relationships, providing a richer characterisation of inter-regional dynamics (Deco et al., 2026).

### Whole-brain modelling in vortex space

The generative model describes the evolution of large-scale brain dynamics within vortex space. It is derived from a system of coupled phase oscillators based on the Kuramoto framework, defined at the level of 1,000 cortical parcels. By applying the Ott–Antonsen ansatz (Ott & Antonsen, 2008), the high-dimensional system can be reduced to a set of coupled Stuart–Landau oscillators, widely used to model brain activity in a supercritical Hopf bifurcation (Deco et al., 2017; Kuznetsov, 2023). Each oscillator thus describes the dynamics of the complex-valued local Kuramoto order parameter of its corresponding macroregion.

In this reduced formulation, each macroregion *σ* in the Schaefer-100 parcellation is modelled as a single complex oscillator *z_σ_*(*t*) ∈ ℂ, whose evolution is governed by:

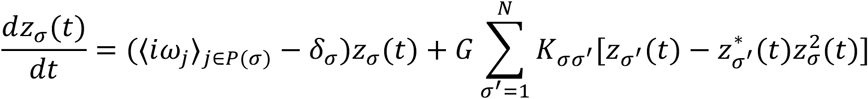

Here, ⟨*iω_j_*⟩_(∈0(*σ*)_denotes the average intrinsic frequency across parcels within macroregion *σ*, estimated from the peak of the empirical BOLD power spectrum using a fast Fourier transform (FFT) and Gaussian smoothing (σ = 0.01 Hz). The damping parameter was fixed at *δ_σ_* = 0.001 for all regions, keeping the system near the oscillatory threshold and supporting metastable dynamics (Ponce-Alvarez & Deco, 2024). Global coupling strength was set to *G* = 0.1, consistent with previous work reproducing resting-state dynamics (Escrichs et al., 2022), while the anatomical coupling matrix K_σσ_′ represents the normalised SC between macroregions. The nonlinear saturation term 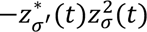 ensures that the trajectories remain bounded and enables the emergence of stable, metastable, and transient dynamics characteristic of resting-state brain activity (Deco et al., 2025).

Simulations were performed using Euler integration with a time step of 0.0625 s. Gaussian noise (*σ* = 0.01) was added to model intrinsic neuronal variability. Each simulation comprised 2,000 time points, with an initial transient discarded for stabilisation, and was repeated 50 times to account for stochastic variability. The resulting simulated time series *z_σ_*(*t*) were used to compute FV following the same procedure as for the empirical data, allowing direct comparison between simulated and observed dynamics.

To infer subject-specific effective vortex interactions, we estimated Generative Effective Connectivity in Vortex Space (GECVO) through an iterative optimisation procedure. The effective coupling matrix was initialised from the empirical structural connectivity, such that 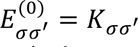. At each iteration, the evolving effective coupling matrix *E*_σσ′_ replaced the anatomical coupling matrix *K*_σσ′_ in the model equations, yielding:

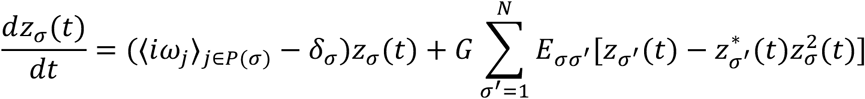

The resulting simulated dynamics were used to compute 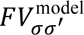, and coupling weights were updated using a pseudo-gradient rule to minimise the discrepancy between empirical and simulated FV:

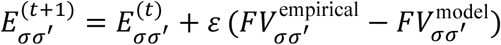

After each update, the coupling matrix was renormalised to preserve consistent scaling across subjects. The learning rate was set to *ε* = 0.01, and optimisation was performed over 1,000 iterations, which was sufficient to obtain a stable and convergent fit. Updates were restricted to existing anatomical connections and inter-hemispheric homotopic pairs, accounting for known limitations of tractography in capturing callosal fibres and spatial resolution (Castaldo et al., 2023).

The converged subject-specific matrix constituted the GECVO, representing the effective coupling strengths required to reproduce the empirical vortex-space dynamics. Model fit was quantified using the Pearson correlation coefficient and mean squared error (MSE) between empirical and simulated FV matrices, which were tracked across optimisation iterations.

### Whole-brain modelling in signal space

To provide a reference framework, we implemented a conventional whole-brain model operating directly in signal space (Deco et al., 2017). Unlike the vortex-space approach, in which local synchrony at the Schaefer-100 level is derived from the fine-grained phase dynamics of constituent Schaefer-1000 parcels, the signal-space model operates directly on BOLD signals averaged within each Schaefer-100 region.

Each brain region was represented as a Stuart–Landau oscillator governed by the normal form of a supercritical Hopf bifurcation. The complex-valued state *z_j_* = *x_j_* + *iy_j_* of region *j* evolved according to:

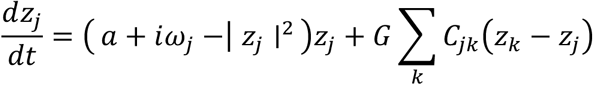

Intrinsic frequencies (*ω_j_*) were estimated from the empirical BOLD power spectrum using the same procedure as in the vortex-space model. The bifurcation parameter was fixed at *a* = −0.01, placing the system just below the transition to self-sustained oscillations. The SC matrix (*C_jk_* in this notation) was globally scaled by a coupling parameter *G* = 0.1, consistent with the previous model.

Numerical simulations used the same integration scheme, time step, noise level, simulation length, transient removal and number of repetitions as the vortex-space model. The real component of each simulated oscillator *z_j_*(*t*) was taken as the synthetic BOLD signal and processed identically to the empirical data, thereby enabling proper comparison.

Model fitting likewise followed the optimisation procedure described for the vortex-space model. Coupling weights were iteratively adjusted to minimise the discrepancy between empirical and simulated signal-space functional connectivity (FC), computed as the Pearson correlation between regional BOLD time series. The coupling matrix C_=>_was then updated according to:

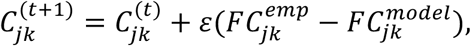

Starting from the SC matrix, coupling weights were iteratively updated according to the previously defined optimisation settings, structural constraints and normalisation procedure. The resulting subject-specific Generative Effective Connectivity (GEC) matrix represents the effective coupling strengths required to reproduce the empirical FC. Model fit was also quantified using the Pearson correlation coefficient and MSE between empirical and simulated FC matrices.

### Machine-learning classification of treatment state and clinical response

Machine learning was used to classify intervention type, treatment phase and clinical response based on subject-specific generative effective connectivity matrices derived from whole-brain modelling. To directly compare the relative sensitivity of signal- and vortex-space representations, all classification analyses were performed in parallel for GEC and GECVO matrices using the same cross-validation, feature-selection and statistical-testing procedures. All matrices were obtained at the level of the Schaefer-100 parcellation and were symmetric by construction.

For each participant and session, the lower triangular elements of each connectivity matrix, excluding the diagonal, were vectorised to yield 4,950 unique effective-connectivity features. Four binary classification tasks were conducted: (i) pre- versus post-treatment psilocybin, (ii) pre- versus post-treatment escitalopram, (iii) psilocybin versus escitalopram based on post-treatment sessions, and (iv) responder versus non-responder classification in the escitalopram group based on pre-treatment connectivity. Response classification was restricted to escitalopram because outcomes were substantially more balanced in this group (8 responders, 12 non-responders) than following psilocybin (18 responders, 4 non-responders).

Classification was implemented using Extreme Gradient Boosting (XGB) with the *scikit-learn* and *xgboost* libraries (Abraham et al., 2014; T. Chen & Guestrin, 2016). XGB was selected for its ability to capture nonlinear interactions while remaining well suited to sparse, high-dimensional feature spaces. This was particularly relevant for neuroimaging datasets, where the number of features greatly exceeds the number of participants and where discriminative information may be distributed across a limited subset of connections. Hyperparameters were fixed a priori and held constant across tasks and connectivity spaces: 200 estimators, a maximum tree depth of 6, a learning rate of 0.1, subsampling of 0.8, column subsampling per tree of 0.8 and log-loss as the evaluation metric. All remaining XGBClassifier parameters were left at their default values, and no hyperparameter tuning was performed.

To prevent information leakage, feature selection was performed exclusively within the training data of each cross-validation fold. The 4,950 edges were ranked according to their statistical discriminability using edge-wise Wilcoxon tests with 10,000 permutations: paired tests for within-subject pre- versus post-treatment classifications and unpaired tests for between-group classifications. Edges were ordered by ascending *P* value, and progressively larger subsets of the highest-ranked features were evaluated using XGB. The feature subset yielding the highest mean cross-validated accuracy was retained for each task and connectivity space. This procedure reduces the influence of weakly informative variables, with the aim of limiting overfitting, improving generalisation, and enhancing classifier performance (R. C. Chen et al., 2020).

Classification performance was evaluated using leave-one-subject-out cross-validation. For longitudinal classifications, one participant was held out at each iteration, with both their pre- and post-treatment sessions excluded from training. For independent-group classifications, one participant from each class was held out at each iteration, and class balance in the training data was enforced by randomly subsampling the larger class to match the smaller class. In both settings, the procedure was repeated over 1,000 random shuffles to obtain stable accuracy estimates. Differences in classification performance between GEC and GECVO were assessed using Wilcoxon tests with 10,000 permutations applied to the corresponding distributions of cross-validated accuracies.

### Regional global brain connectivity

For each participant and session, regional global brain connectivity (GBC) was computed as the mean effective connectivity of each Schaefer-100 region with the rest of the brain. GBC was calculated independently from GECVO and GEC matrices in vortex and signal spaces, respectively. Treatment-related changes were quantified as post-treatment minus pre-treatment GBC and assessed separately within each intervention using paired Wilcoxon tests and between interventions using unpaired Wilcoxon tests, with 10,000 permutations in each case. *P* values were corrected across regions using FDR, and effect sizes were quantified using Cohen’s *d*.

### Connectome-based modelling of clinical improvement

To further examine the relationship between baseline effective connectivity and subsequent clinical improvement, we applied a linear association framework adapted from connectome-based predictive modelling (Shen et al., 2017). Signal-space (GEC) and vortex-space (GECVO) effective-connectivity matrices were analysed separately to quantify descriptive in-sample associations with subsequent clinical improvement and identify the contributing connectivity features.

Baseline regional GBC values, computed as described above, were used as predictors of subsequent clinical improvement. Improvement was defined as the pre-treatment minus post-treatment change in BDI score, such that larger positive values indicated greater symptom reduction.

Within each representation, regional GBC was correlated with clinical improvement across participants. At correlation thresholds ranging from |r| = 0.10 to 0.60 in increments of 0.02, regions showing positive and negative associations were identified separately. Participant-specific positive and negative network scores were then calculated by summing GBC values across the corresponding sets of regions. These scores were entered as predictors into an ordinary least-squares linear model with an intercept, fitted in MATLAB using the *fitlm* function. No regularisation was applied.

Model fit was evaluated at each threshold using the correlation between observed and model-fitted clinical improvement, the ordinary R², and the overall model P value. Overall model significance was assessed using the F-test implemented in *coefTest*. The threshold yielding the highest in-sample fitted correlation was retained for each connectivity space, together with the numbers of positively and negatively associated regions entering the model.

### Network modularity of effective connectivity

To assess treatment-related changes in network segregation, we computed the modularity of subject-specific effective connectivity matrices in both signal space (GEC) and vortex space (GECVO). Modularity quantifies the extent to which connectivity is concentrated within communities relative to that expected under a strength-preserving null model. For each participant and session, self-connections were excluded, and modularity was evaluated using the Newman–Girvan quality function (Newman & Girvan, 2004):

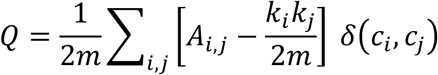

where *A_i,j_* is effective coupling between regions *i* and *j*, *k_i_* and *k*_j_ denote the corresponding nodal strengths, *m* is the total connection weight in the network, and *δ*(*c_i_*, *c_j_*) is 1 when nodes *i* and *j* are assigned to the same community and 0 otherwise.

Community structure was estimated using the Louvain algorithm (Blondel et al., 2008) with resolution parameter γ = 1. Given the stochastic nature of Louvain optimisation, the algorithm was repeated 100 times for each subject and session, and the partition yielding the highest modularity value was retained. The modularity value from this optimal partition was used for subsequent analyses, conducted separately for psilocybin and escitalopram before and after treatment. Within each representation, pre–post changes within treatments and post-minus-pre differences between treatments were assessed using paired and unpaired Wilcoxon tests, respectively, with 10,000 permutations.

### Statistical analyses

Statistical comparisons were performed using paired or unpaired permutation-based Wilcoxon tests (10,000 permutations), as appropriate, with effect sizes quantified using Cohen’s *d* where applicable. Associations were assessed using Pearson correlations. Multiple comparisons across brain regions were corrected using the Benjamini–Hochberg false discovery rate (FDR) procedure (Benjamini & Hochberg, 1995). Statistical significance was set at *P* < 0.05.

## Declarations

### Author contributions

Álvaro Acosta Gordo: Conceptualization; Data curation; Formal analysis; Investigation; Methodology; Software; Validation; Visualization; Writing—original draft; Writing—review & editing. Paulina Clara Dagnino: Conceptualization; Data curation; Methodology; Software; Supervision; Writing—review & editing. Yonatan Sanz Perl: Software; Data curation; Writing—review & editing. Robin L. Carhart-Harris: Data curation; Funding acquisition; Project administration; Resources; Writing—review & editing. David Erritzoe: Data curation; Funding acquisition; Project administration; Resources; Writing—review & editing. David J. Nutt: Data curation; Funding acquisition; Project administration; Resources; Writing—review & editing. Morten L. Kringelbach: Data curation; Funding acquisition; Project administration; Resources; Supervision; Writing—review & editing. Gustavo Deco: Conceptualization; Data curation; Funding acquisition; Methodology; Project administration; Resources; Software; Supervision; Writing—review & editing.

### Data availability

Requests for access to raw and processed participant-level neuroimaging and clinical data from the clinical trial are reviewed by R.L.C.-H., the chief investigator of the original work, and are subject to participant consent, confidentiality requirements and applicable data-sharing agreements. The group-average structural connectivity was derived from diffusion MRI data from the Human Connectome Project; access to the source data is available through the HCP subject to its data-use terms.

### Code availability

The analysis code used in this study is publicly available in the GitHub repository: https://github.com/alvaroacostagordo/vortex-wbm-psilocybin-escitalopram.

## Acknowledgements

G.D. was supported by Grant PID2022-136216NB-I00 funded by MICIU/AEI/10.13039/501100011033 and by the European Regional Development Fund (ERDF, “A way of making Europe”); by the EU-funded ERC Synergy Horizon Europe project NEurological MEchanismS of Injury, and Sleep-like cellular dynamics (NEMESIS; ref. 101071900); by the AGAUR research support grant (ref. 2021 SGR 00917) funded by the Department of Research and Universities of the Generalitat de Catalunya; and by Grant PID2024-155136NI-I00 funded by MICIU/AEI/10.13039/501100011033 and by the European Regional Development Fund (ERDF, “A way of making Europe”).

P.C.D. was supported by the Department of Research and Universities of the Generalitat de Catalunya through the Agency for Management of University and Research Grants (AGAUR) FI-SDUR programme (Grant No. 2022 FISDU 00229).

Y.S.P. was supported by the EU-funded ERC Synergy Horizon Europe project NEurological MEchanismS of Injury, and Sleep-like cellular dynamics (NEMESIS; ref. 101071900), and by Grant PID2024-162576NA-I00 funded by MICIU/AEI/10.13039/501100011033 and by the European Regional Development Fund (ERDF, “A way of making Europe”).

M.L.K. was supported by the Centre for Eudaimonia and Human Flourishing, funded by the Pettit and Carlsberg Foundations, and the Center for Music in the Brain, funded by the Danish National Research Foundation (DNRF117).

Funding for the double-blind randomised controlled trial was provided by a private donation from the Alexander Mosley Charitable Trust, with additional support from the founding partners of Imperial College London’s Centre for Psychedelic Research. The research was conducted at the National Institute for Health and Care Research (NIHR) Imperial Clinical Research Facility (ICRF). The funding bodies had no role in the study design, data collection and analysis, decision to publish, or preparation of the manuscript.

## References

Abraham, A., Pedregosa, F., Eickenberg, M., Gervais, P., Mueller, A., Kossaifi, J., Gramfort, A., Thirion, B., & Varoquaux, G. (2014). Machine learning for neuroimaging with scikit-learn. Frontiers in Neuroinformatics, 8, 14. 10.3389/fninf.2014.00014

Avants, B., Tustison, N. J., & Song, G. (2009). Advanced Normalization Tools: V1.0. The Insight Journal. 10.54294/UVNHIN

Beck, A. T., Ward, C. H., Mendelson, M., Mock, J., & Erbaugh, J. (1961). An Inventory for Measuring Depression. Archives of General Psychiatry, 4(6), 561–571. 10.1001/ARCHPSYC.1961.01710120031004

Benjamini, Y., & Hochberg, Y. (1995). Controlling the False Discovery Rate: A Practical and Powerful Approach to Multiple Testing. Journal of the Royal Statistical Society Series B: Statistical Methodology, 57(1), 289–300. 10.1111/J.2517-6161.1995.TB02031.X

Blondel, V. D., Guillaume, J. L., Lambiotte, R., & Lefebvre, E. (2008). Fast unfolding of communities in large networks. Journal of Statistical Mechanics: Theory and Experiment, 2008(10), P10008. 10.1088/1742-5468/2008/10/p10008

Carhart-Harris, R. L., Giribaldi, B., Watts, R., Baker-Jones, M., Murphy-Beiner, A., Murphy, R., Martell, J., Blemings, A., Erritzoe, D., & Nutt, D. J. (2021). Trial of Psilocybin versus Escitalopram for Depression. New England Journal of Medicine, 384(15), 1402–1411. 10.1056/NEJMoa2032994

Castaldo, F., Páscoa dos Santos, F., Timms, R. C., Cabral, J., Vohryzek, J., Deco, G., Woolrich, M., Friston, K., Verschure, P., & Litvak, V. (2023). Multi-modal and multi-model interrogation of large-scale functional brain networks. NeuroImage, 277, 120236. 10.1016/J.NEUROIMAGE.2023.120236

Chen, R. C., Dewi, C., Huang, S. W., & Caraka, R. E. (2020). Selecting critical features for data classification based on machine learning methods. Journal of Big Data, 7(1), 52. 10.1186/s40537-020-00327-4

Chen, T., & Guestrin, C. (2016). XGBoost: A scalable tree boosting system. Proceedings of the 22nd ACM SIGKDD International Conference on Knowledge Discovery and Data Mining, 13-17-August-2016, 785–794. 10.1145/2939672.2939785

Cipriani, A., Furukawa, T. A., Salanti, G., Chaimani, A., Atkinson, L. Z., Ogawa, Y., Leucht, S., Ruhe, H. G., Turner, E. H., Higgins, J. P. T., Egger, M., Takeshima, N., Hayasaka, Y., Imai, H., Shinohara, K., Tajika, A., Ioannidis, J. P. A., & Geddes, J. R. (2018). Comparative efficacy and acceptability of 21 antidepressant drugs for the acute treatment of adults with major depressive disorder: a systematic review and network meta-analysis. The Lancet, 391(10128), 1357–1366. 10.1016/S0140-6736(17)32802-7

Cox, R. W. (1996). AFNI: Software for Analysis and Visualization of Functional Magnetic Resonance Neuroimages. Computers and Biomedical Research, 29(3), 162–173. 10.1006/CBMR.1996.0014

Cruzat, J., Sanz Perl, Y., Escrichs, A., Vohryzek, J., Timmermann, C., Roseman, L., Luppi, A. I., Ibañez, A., Nutt, D., Carhart-Harris, R. L., Tagliazucchi, E., Deco, G., & Kringelbach, M. L. (2022). Effects of classic psychedelic drugs on turbulent signatures in brain dynamics. Network Neuroscience, 6(4), 1104–1124. 10.1162/NETN_A_00250

Dagnino, P. C., Acero-Pousa, I., Zamora-Lopez, G., Escrichs, A., Erritzoe, D., Nutt, D. J., Carhart-Harris, R. L., Sanz Perl, Y., Kringelbach, M. L., & Deco, G. (2026). Distinct brain responses to psilocybin and escitalopram in depression captured by the Fluctuation-Dissipation Theorem. 10.64898/2026.06.12.731811

Dale, A. M., Fischl, B., & Sereno, M. I. (1999). Cortical Surface-Based Analysis: I. Segmentation and Surface Reconstruction. NeuroImage, 9(2), 179–194. 10.1006/NIMG.1998.0395

Daws, R. E., Timmermann, C., Giribaldi, B., Sexton, J. D., Wall, M. B., Erritzoe, D., Roseman, L., Nutt, D., & Carhart-Harris, R. L. (2022). Increased global integration in the brain after psilocybin therapy for depression. Nature Medicine, 28(4), 844–851. 10.1038/S41591-022-01744-Z

Deco, G., & Kringelbach, M. L. (2014). Great expectations: Using whole-brain computational connectomics for understanding neuropsychiatric disorders. Neuron, 84(5), 892–905. 10.1016/j.neuron.2014.08.034

Deco, G., & Kringelbach, M. L. (2020). Turbulent-like Dynamics in the Human Brain. Cell Reports, 33(10), 108471. 10.1016/j.celrep.2020.108471

Deco, G., Kringelbach, M. L., Jirsa, V. K., & Ritter, P. (2017). The dynamics of resting fluctuations in the brain: metastability and its dynamical cortical core. Scientific Reports, 7(1), 3095. 10.1038/s41598-017-03073-5

Deco, G., Liebana Garcia, S., Sanz Perl, Y., Sporns, O., & Kringelbach, M. L. (2023). The effect of turbulence in brain dynamics information transfer measured with magnetoencephalography. Communications Physics, 6(1), 74. 10.1038/s42005-023-01192-2

Deco, G., Sanz Perl, Y., Feng, J., & Kringelbach, M. L. (2026). The turbulent brain: Modeling vortex interactions for understanding human cognition. Network Neuroscience, 10(3), 630–654. 10.1162/NETN.A.554

Deco, G., Sanz Perl, Y., Jerotic, K., Escrichs, A., & Kringelbach, M. L. (2025). Turbulence as a framework for brain dynamics in health and disease. Neuroscience & Biobehavioral Reviews, 169, 105988. 10.1016/J.NEUBIOREV.2024.105988

Deco, G., Sanz Perl, Y., Johnson, S., Bourke, N., Carhart-Harris, R. L., & Kringelbach, M. L. (2024). Different hierarchical reconfigurations in the brain by psilocybin and escitalopram for depression. Nature Mental Health, 2(9), 1096–1110. 10.1038/s44220-024-00298-y

DeMaster, D., Godlewska, B., Liang, M., Vannucci, M., Bockmann, T., Bo, C., & Selvaraj, S. (2022). Effective connectivity between resting-state networks in depression. Journal of Affective Disorders, 307, 79–86. 10.1016/J.JAD.2022.03.041

Escrichs, A., Sanz Perl, Y., Fisher, P. M., Martínez-Molina, N., G-Guzman, E., Frokjaer, V. G., Kringelbach, M. L., Knudsen, G. M., & Deco, G. (2025). Whole-brain turbulent dynamics predict responsiveness to pharmacological treatment in major depressive disorder. Molecular Psychiatry, 30(3), 1069–1079. 10.1038/s41380-024-02690-7

Escrichs, A., Sanz Perl, Y., Uribe, C., Camara, E., Türker, B., Pyatigorskaya, N., López-González, A., Pallavicini, C., Panda, R., Annen, J., Gosseries, O., Laureys, S., Naccache, L., Sitt, J. D., Laufs, H., Tagliazucchi, E., Kringelbach, M. L., & Deco, G. (2022). Unifying turbulent dynamics framework distinguishes different brain states. Communications Biology, 5(1), 1–13. 10.1038/s42003-022-03576-6

Fox, M. D., Snyder, A. Z., Vincent, J. L., Corbetta, M., Van Essen, D. C., & Raichle, M. E. (2005). The human brain is intrinsically organized into dynamic, anticorrelated functional networks. Proceedings of the National Academy of Sciences of the United States of America, 102(27), 9673–9678. 10.1073/pnas.0504136102

Girn, M., Doss, M. K., Roseman, L., Preller, K. H., Palhano-Fontes, F., Pasquini, L., Barrett, F. S., Mallaroni, P., Mason, N. L., Timmermann, C., McCulloch, D. E., Fisher, P. M., Winston, B. S., Moujaes, F., Muller, F., Liechti, M. E., Vollenweider, F. X., Ramaekers, J. G., Kuypers, K., … Bzdok, D. (2026). An international mega-analysis of psychedelic drug effects on brain circuit function. Nature Medicine, 32(4), 1543–1554. 10.1038/s41591-026-04287-9

Hancock, F., Cabral, J., Luppi, A. I., Rosas, F. E., Mediano, P. A. M., Dipasquale, O., & Turkheimer, F. E. (2022). Metastability, fractal scaling, and synergistic information processing: What phase relationships reveal about intrinsic brain activity. NeuroImage, 259, 119433. 10.1016/J.NEUROIMAGE.2022.119433

Horn, A., & Blankenburg, F. (2016). Toward a standardized structural–functional group connectome in MNI space. NeuroImage, 124, 310–322. 10.1016/J.NEUROIMAGE.2015.08.048

Horn, A., Neumann, W. J., Degen, K., Schneider, G. H., & Kühn, A. A. (2017). Toward an electrophysiological “sweet spot” for deep brain stimulation in the subthalamic nucleus. Human Brain Mapping, 38(7), 3377–3390. 10.1002/HBM.23594

Kawamura, Y., Nakao, H., & Kuramoto, Y. (2007). Noise-induced turbulence in nonlocally coupled oscillators. Physical Review E, 75(3), 036209. 10.1103/PHYSREVE.75.036209

Korgaonkar, M. S., Goldstein-Piekarski, A. N., Fornito, A., & Williams, L. M. (2019). Intrinsic connectomes are a predictive biomarker of remission in major depressive disorder. Molecular Psychiatry, 25(7), 1537–1549. 10.1038/s41380-019-0574-2

Kringelbach, M. L., Cruzat, J., Cabral, J., Knudsen, G. M., Carhart-Harris, R. L., Whybrow, P. C., Logothetis, N. K., & Deco, G. (2020). Dynamic coupling of whole-brain neuronal and neurotransmitter systems. Proceedings of the National Academy of Sciences of the United States of America, 117(17), 9566–9576. 10.1073/PNAS.1921475117

Kuramoto, Y. (1984). Chemical Oscillations, Waves, and Turbulence (Vol. 19). Springer. 10.1007/978-3-642-69689-3

Kuznetsov, Y. A. (2023). Elements of Applied Bifurcation Theory (4th ed., Vol. 112). Springer. 10.1007/978-3-031-22007-4

Landy, K., Rosani, A., & Estevez, R. (2023). Escitalopram. In *StatPearls*. StatPearls Publishing. https://www.ncbi.nlm.nih.gov/books/NBK557734/

Lowe, M. J., Mock, B. J., & Sorenson, J. A. (1998). Functional connectivity in single and multislice echoplanar imaging using resting-state fluctuations. NeuroImage, 7(2), 119–132. 10.1006/NIMG.1997.0315

Lundberg, J., Christophersen, J. S., Petersen, K. B., Loft, H., Halldin, C., & Farde, L. (2007). PET measurement of serotonin transporter occupancy: a comparison of escitalopram and citalopram. International Journal of Neuropsychopharmacology, 10(6), 777–785. 10.1017/S1461145706007486

Madsen, M. K., Fisher, P. M., Burmester, D., Dyssegaard, A., Stenbæk, D. S., Kristiansen, S., Johansen, S. S., Lehel, S., Linnet, K., Svarer, C., Erritzoe, D., Ozenne, B., & Knudsen, G. M. (2019). Psychedelic effects of psilocybin correlate with serotonin 2A receptor occupancy and plasma psilocin levels. Neuropsychopharmacology, 44(7), 1328–1334. 10.1038/s41386-019-0324-9

Maier-Hein, K. H., Neher, P. F., Houde, J. C., Côté, M. A., Garyfallidis, E., Zhong, J., Chamberland, M., Yeh, F. C., Lin, Y. C., Ji, Q., Reddick, W. E., Glass, J. O., Chen, D. Q., Feng, Y., Gao, C., Wu, Y., Ma, J., Renjie, H., Li, Q., … Descoteaux, M. (2017). The challenge of mapping the human connectome based on diffusion tractography. Nature Communications, 8(1), 1349. 10.1038/s41467-017-01285-x

Mnookin, S. (2016). Out of the shadows: making mental health a global development priority. https://documents.worldbank.org/en/publication/documents-reports/documentdetail/270131468187759113/out-of-the-shadows-making-mental-health-a-global-development-priority

Newman, M. E. J., & Girvan, M. (2004). Finding and evaluating community structure in networks. Physical Review E, 69(2), 026113. 10.1103/PhysRevE.69.026113

Nichols, D. E. (2016). Psychedelics. Pharmacological Reviews, 68(2), 264–355. 10.1124/PR.115.011478

Nunez, J. J., Liu, Y. S., Cao, B., Frey, B. N., Ho, K., Milev, R., Müller, D. J., Rotzinger, S., Soares, C. N., Taylor, V. H., Uher, R., Kennedy, S. H., & Lam, R. W. (2023). Response trajectories during escitalopram treatment of patients with major depressive disorder. Psychiatry Research, 327, 115361. 10.1016/J.PSYCHRES.2023.115361

Ott, E., & Antonsen, T. M. (2008). Low dimensional behavior of large systems of globally coupled oscillators. Chaos, 18(3), 037113. 10.1063/1.2930766

Otte, C., Gold, S. M., Penninx, B. W., Pariante, C. M., Etkin, A., Fava, M., Mohr, D. C., & Schatzberg, A. F. (2016). Major depressive disorder. Nature Reviews. Disease Primers, 2, 16065. 10.1038/NRDP.2016.65

Patow, G., Martin, I., Sanz Perl, Y., Kringelbach, M. L., & Deco, G. (2024). Whole-brain modelling: an essential tool for understanding brain dynamics. Nature Reviews Methods Primers, 4(1), 1–2. 10.1038/s43586-024-00336-0

Ponce-Alvarez, A., & Deco, G. (2024). The Hopf whole-brain model and its linear approximation. Scientific Reports, 14(1), 1–12. 10.1038/s41598-024-53105-0

Rucker, J. J. (2024). Evidence versus expectancy: the development of psilocybin therapy. BJPsych Bulletin, 48(2), 110–117. 10.1192/BJB.2023.28

Rush, A. J., Trivedi, M. H., Wisniewski, S. R., Nierenberg, A. A., Stewart, J. W., Warden, D., Niederehe, G., Thase, M. E., Lavori, P. W., Lebowitz, B. D., McGrath, P. J., Rosenbaum, J. F., Sackeim, H. A., Kupfer, D. J., Luther, J., & Fava, M. (2006). Acute and longer-term outcomes in depressed outpatients requiring one or several treatment steps: A STAR*D report. American Journal of Psychiatry, 163(11), 1905–1917. 10.1176/ajp.2006.163.11.1905

Sanz Perl, Y., Mininni, P., Tagliazucchi, E., Kringelbach, M. L., & Deco, G. (2023). Scaling of whole-brain dynamics reproduced by high-order moments of turbulence indicators. Physical Review Research, 5(3), 033183. 10.1103/PhysRevResearch.5.033183

Sanz Perl, Y., Zamora-Lopez, G., Montbrió, E., Monge-Asensio, M., Vohryzek, J., Fittipaldi, S., Campo, C. G., Moguilner, S., Ibañez, A., Tagliazucchi, E., Yeo, B. T. T., Kringelbach, M. L., & Deco, G. (2023). The impact of regional heterogeneity in whole-brain dynamics in the presence of oscillations. Network Neuroscience, 7(2), 632–660. 10.1162/NETN_A_00299

Schaefer, A., Kong, R., Gordon, E. M., Laumann, T. O., Zuo, X.-N., Holmes, A. J., Eickhoff, S. B., & Yeo, B. T. T. (2018). Local-Global Parcellation of the Human Cerebral Cortex from Intrinsic Functional Connectivity MRI. Cerebral Cortex, 28(9), 3095–3114. 10.1093/CERCOR/BHX179

Setsompop, K., Kimmlingen, R., Eberlein, E., Witzel, T., Cohen-Adad, J., McNab, J. A., Keil, B., Tisdall, M. D., Hoecht, P., Dietz, P., Cauley, S. F., Tountcheva, V., Matschl, V., Lenz, V. H., Heberlein, K., Potthast, A., Thein, H., Van Horn, J., Toga, A., … Wald, L. L. (2013). Pushing the limits of in vivo diffusion MRI for the Human Connectome Project. NeuroImage, 80, 220– 233. 10.1016/J.NEUROIMAGE.2013.05.078

Shen, X., Finn, E. S., Scheinost, D., Rosenberg, M. D., Chun, M. M., Papademetris, X., & Constable, R. T. (2017). Using connectome-based predictive modeling to predict individual behavior from brain connectivity. Nature Protocols, 12(3), 506–518. 10.1038/nprot.2016.178

Smith, S. M., Jenkinson, M., Woolrich, M. W., Beckmann, C. F., Behrens, T. E. J., Johansen-Berg, H., Bannister, P. R., De Luca, M., Drobnjak, I., Flitney, D. E., Niazy, R. K., Saunders, J., Vickers, J., Zhang, Y., De Stefano, N., Brady, J. M., & Matthews, P. M. (2004). Advances in functional and structural MR image analysis and implementation as FSL. NeuroImage, 23(SUPPL. 1), S208–S219. 10.1016/J.NEUROIMAGE.2004.07.051

Socoró-Garrigosa, M., Sanz Perl, Y., Kringelbach, M. L., Erritzoe, D., Nutt, D. J., Carhart-Harris, R. L., Vohryzek, J., & Deco, G. (2025). Perturbing whole-brain models of brain hierarchy: An application for depression following pharmacological treatment. Annals of the New York Academy of Sciences, 1550(1), 255–272. 10.1111/NYAS.15391

Tagliazucchi, E., Roseman, L., Kaelen, M., Orban, C., Muthukumaraswamy, S. D., Murphy, K., Laufs, H., Leech, R., McGonigle, J., Crossley, N., Bullmore, E., Williams, T., Bolstridge, M., Feilding, A., Nutt, D. J., & Carhart-Harris, R. (2016). Increased Global Functional Connectivity Correlates with LSD-Induced Ego Dissolution. Current Biology, 26(8), 1043–1050. 10.1016/J.CUB.2016.02.010

Vohryzek, J., Cabral, J., Lord, L. D., Fernandes, H. M., Roseman, L., Nutt, D. J., Carhart-Harris, R. L., Deco, G., & Kringelbach, M. L. (2024). Brain dynamics predictive of response to psilocybin for treatment-resistant depression. Brain Communications, 6(2). 10.1093/braincomms/fcae049

Wang, Y., Zhou, J., Chen, X., Liu, R., Zhang, Z., Feng, L., Feng, Y., Wang, G., & Zhou, Y. (2024). Effects of escitalopram therapy on effective connectivity among core brain networks in major depressive disorder. Journal of Affective Disorders, 350, 39–48. 10.1016/J.JAD.2024.01.115

World Health Organization. (2025, August 29). Depressive disorder (depression). https://www.who.int/news-room/fact-sheets/detail/depression

Yeo, B. T. T., Krienen, F. M., Sepulcre, J., Sabuncu, M. R., Lashkari, D., Hollinshead, M., Roffman, J. L., Smoller, J. W., Zöllei, L., Polimeni, J. R., Fisch, B., Liu, H., & Buckner, R. L. (2011). The organization of the human cerebral cortex estimated by intrinsic functional connectivity. Journal of Neurophysiology, 106(3), 1125–1165. 10.1152/jn.00338.2011

