## Supplementary Tables for "Modelling vortex interactions reveals distinct patterns of brain reconfiguration following psilocybin and escitalopram"

| Model | Before psilocybin | After psilocybin | Before escitalopram | After escitalopram |
|---|---|---|---|---|
| Vortex space | 0.6429 ± 0.0453 | 0.6298 ± 0.0400 | 0.6285 ± 0.0423 | 0.6125 ± 0.0417 |
| Signal space | 0.7484 ± 0.0382 | 0.7555 ± 0.0434 | 0.7310 ± 0.0496 | 0.7481 ± 0.0310 |

**Supplementary Table S1. Whole-brain model fit across treatments and sessions.** Mean ± s.d. of the Pearson correlation between empirical and simulated observables for vortex-space and signal-space whole-brain models across subjects and experimental conditions.

| Model | Before psilocybin | After psilocybin | Before escitalopram | After escitalopram |
|---|---|---|---|---|
| Vortex space | 0.0082 ± 0.0013 | 0.0071 ± 0.0010 | 0.0076 ± 0.0016 | 0.0077 ± 0.0011 |
| Signal space | 0.0858 ± 0.0355 | 0.0712 ± 0.0258 | 0.0662 ± 0.0212 | 0.0726 ± 0.0249 |

**Supplementary Table S2. Whole-brain model reconstruction error across treatments and sessions.** Mean ± s.d. of the reconstruction error (MSE) for vortex-space and signal-space whole-brain models across subjects and experimental conditions.

| Yeo network | No. of regions | Psilocybin, n (%) | Escitalopram, n (%) | Between treatments, n (%) |
|---|---|---|---|---|
| Visual | 17 | 13 (76.5%) | 0 (0.0%) | 12 (70.6%) |
| Somatomotor | 14 | 13 (92.9%) | 0 (0.0%) | 8 (57.1%) |
| Dorsal attention | 15 | 14 (93.3%) | 0 (0.0%) | 12 (80.0%) |
| Salience / ventral attention | 12 | 9 (75.0%) | 0 (0.0%) | 3 (25.0%) |
| Limbic | 5 | 3 (60.0%) | 0 (0.0%) | 3 (60.0%) |
| Control | 13 | 7 (53.8%) | 0 (0.0%) | 9 (69.2%) |
| Default mode | 24 | 17 (70.8%) | 0 (0.0%) | 17 (70.8%) |
| Total | 100 | 76 (76.0%) | 0 (0.0%) | 64 (64.0%) |

**Supplementary Table S3. Distribution of significant regional GBC effects across canonical Yeo networks in vortex space.** Values indicate the number and percentage of Schaefer-100 regions within each network surviving FDR correction ($q < 0.05$) for the psilocybin and escitalopram pre–post comparisons and the between-treatment comparison. Percentages are relative to the total number of regions assigned to each network.

| Yeo network | No. of regions | Psilocybin, n (%) | Escitalopram, n (%) | Between treatments, n (%) |
|---|---|---|---|---|
| Visual | 17 | 0 (0.0%) | 0 (0.0%) | 0 (0.0%) |
| Somatomotor | 14 | 0 (0.0%) | 0 (0.0%) | 0 (0.0%) |
| Dorsal attention | 15 | 0 (0.0%) | 0 (0.0%) | 0 (0.0%) |
| Salience / ventral attention | 12 | 0 (0.0%) | 0 (0.0%) | 0 (0.0%) |
| Limbic | 5 | 0 (0.0%) | 0 (0.0%) | 0 (0.0%) |
| Control | 13 | 0 (0.0%) | 0 (0.0%) | 0 (0.0%) |
| Default mode | 24 | 0 (0.0%) | 0 (0.0%) | 0 (0.0%) |
| Total | 100 | 0 (0.0%) | 0 (0.0%) | 0 (0.0%) |

**Supplementary Table S4. Distribution of significant regional GBC effects across canonical Yeo networks in signal space.** Values indicate the number and percentage of Schaefer-100 regions within each network surviving FDR correction ($q < 0.05$) for the psilocybin and escitalopram pre–post comparisons and the between-treatment comparison. Percentages are relative to the total number of regions assigned to each network.

| Region | Yeo network | Mean ΔGBC | Cohen's d | Permutation p-value | FDR q-value |
|---|---|---|---|---|---|
| Left visual 9 | Visual | 0.0158 | 0.90 | 0.0006 | 0.0030 |
| Left dorsal attention posterior 1 | Dorsal attention | 0.0156 | 1.26 | 0.0001 | 0.0024 |
| Left dorsal attention posterior 5 | Dorsal attention | 0.0155 | 0.94 | 0.0003 | 0.0024 |
| Left somatomotor 3 | Somatomotor | 0.0142 | 1.01 | 0.0003 | 0.0024 |
| Right dorsal attention posterior 4 | Dorsal attention | 0.0138 | 0.91 | 0.0003 | 0.0024 |
| Right dorsal attention posterior 1 | Dorsal attention | 0.0135 | 0.88 | 0.0010 | 0.0039 |
| Right somatomotor 7 | Somatomotor | 0.0130 | 0.71 | 0.0037 | 0.0080 |
| Right dorsal attention posterior 3 | Dorsal attention | 0.0128 | 0.84 | 0.0003 | 0.0024 |
| Right dorsal attention posterior 5 | Dorsal attention | 0.0127 | 1.20 | 0.0002 | 0.0024 |
| Right somatomotor 2 | Somatomotor | 0.0125 | 0.71 | 0.0084 | 0.0153 |
| Right cingulate 1 | Control | 0.0123 | 1.12 | 0.0004 | 0.0024 |
| Right dorsal attention posterior 2 | Dorsal attention | 0.0113 | 1.15 | 0.0003 | 0.0024 |
| Right dorsal attention precentral ventral 1 | Dorsal attention | 0.0113 | 0.80 | 0.0006 | 0.0030 |
| Right visual 7 | Visual | 0.0113 | 0.89 | 0.0013 | 0.0039 |
| Left cingulate 1 | Control | 0.0112 | 0.67 | 0.0069 | 0.0133 |
| Right temporo-occipito-parietal junction 2 | Salience / ventral attention | 0.0111 | 0.72 | 0.0013 | 0.0039 |
| Left somatomotor 1 | Somatomotor | 0.0111 | 0.95 | 0.0004 | 0.0024 |
| Right somatomotor 5 | Somatomotor | 0.0110 | 0.91 | 0.0003 | 0.0024 |
| Right visual 5 | Visual | 0.0108 | 0.88 | 0.0003 | 0.0024 |
| Right visual 8 | Visual | 0.0106 | 0.82 | 0.0027 | 0.0066 |

**Supplementary Table S5. Regions showing the largest treatment-related changes in vortex-space regional GBC following psilocybin.** The top 20% of Schaefer-100 regions are ranked by |mean ΔGBC|, where ΔGBC denotes the post- minus pre-treatment change. p-values and FDR-corrected q-values correspond to the regional tests across all 100 regions.

| Region | Yeo network | Mean ΔGBC | Cohen’s d | Permutation p-value | FDR q-value |
|---|---|---|---|---|---|
| Left visual 9 | Visual | -0.0072 | -0.56 | 0.0738 | 0.4940 |
| Right dorsal attention posterior 3 | Dorsal attention | -0.0071 | -0.44 | 0.1016 | 0.5976 |
| Left precuneus 1 | Control | -0.0065 | -0.49 | 0.0488 | 0.4940 |
| Right somatomotor 2 | Somatomotor | -0.0063 | -0.48 | 0.0438 | 0.4940 |
| Right visual 6 | Visual | -0.0057 | -0.57 | 0.0267 | 0.4940 |
| Left prefrontal cortex 2 | Default mode | -0.0056 | -0.57 | 0.0257 | 0.4940 |
| Right dorsal attention posterior 2 | Dorsal attention | -0.0055 | -0.49 | 0.0527 | 0.4940 |
| Right dorsal/medial prefrontal cortex 2 | Default mode | -0.0055 | -0.50 | 0.0498 | 0.4940 |
| Left cingulate 1 | Control | -0.0052 | -0.31 | 0.1948 | 0.6299 |
| Left dorsal attention posterior 6 | Dorsal attention | -0.0051 | -0.38 | 0.0741 | 0.4940 |
| Right dorsal attention frontal eye field 1 | Dorsal attention | -0.0051 | -0.52 | 0.0643 | 0.4940 |
| Left lateral prefrontal cortex 1 | Control | -0.0051 | -0.66 | 0.0104 | 0.4940 |
| Right medial posterior prefrontal cortex 1 | Control | -0.0050 | -0.38 | 0.0381 | 0.4940 |
| Left lateral prefrontal cortex 1 | Salience / ventral attention | -0.0048 | -0.28 | 0.2842 | 0.6315 |
| Left prefrontal cortex 3 | Default mode | -0.0045 | -0.40 | 0.1248 | 0.6299 |
| Right medial 1 | Salience / ventral attention | -0.0044 | -0.36 | 0.1458 | 0.6299 |
| Right visual 8 | Visual | -0.0044 | -0.42 | 0.0640 | 0.4940 |
| Left somatomotor 5 | Somatomotor | -0.0044 | -0.33 | 0.1833 | 0.6299 |
| Left prefrontal cortex 6 | Default mode | -0.0044 | -0.32 | 0.1783 | 0.6299 |
| Left dorsal attention posterior 1 | Dorsal attention | 0.0043 | 0.25 | 0.1825 | 0.6299 |

**Supplementary Table S6. Regions showing the largest treatment-related changes in vortex-space regional GBC following escitalopram.** The top 20% of Schaefer-100 regions are ranked by |mean ΔGBC|, where ΔGBC denotes the post- minus pre-treatment change. p-values and FDR-corrected q-values correspond to the regional tests across all 100 regions.

| Region | Yeo network | Treatment difference | Cohen's d | Permutation p-value | FDR q-value |
|---|---|---|---|---|---|
| Left visual 9 | Visual | 0.0230 | 1.49 | 0.0001 | 0.0033 |
| Right dorsal attention posterior 3 | Dorsal attention | 0.0198 | 1.27 | 0.0001 | 0.0033 |
| Right somatomotor 2 | Somatomotor | 0.0188 | 1.20 | 0.0009 | 0.0045 |
| Right dorsal attention posterior 4 | Dorsal attention | 0.0179 | 1.20 | 0.0006 | 0.0037 |
| Left dorsal attention posterior 5 | Dorsal attention | 0.0168 | 1.10 | 0.0021 | 0.0066 |
| Right dorsal attention posterior 2 | Dorsal attention | 0.0168 | 1.61 | 0.0001 | 0.0033 |
| Left cingulate 1 | Control | 0.0164 | 0.98 | 0.0020 | 0.0065 |
| Right visual 6 | Visual | 0.0152 | 1.38 | 0.0004 | 0.0037 |
| Right visual 8 | Visual | 0.0150 | 1.27 | 0.0005 | 0.0037 |
| Right medial 1 | Salience / ventral attention | 0.0150 | 1.16 | 0.0005 | 0.0037 |
| Left lateral prefrontal cortex 1 | Salience / ventral attention | 0.0149 | 0.87 | 0.0113 | 0.0237 |
| Right dorsal/medial prefrontal cortex 2 | Default mode | 0.0148 | 1.32 | 0.0003 | 0.0037 |
| Left dorsal attention posterior 6 | Dorsal attention | 0.0148 | 1.04 | 0.0006 | 0.0037 |
| Right dorsal attention posterior 5 | Dorsal attention | 0.0147 | 1.42 | 0.0002 | 0.0037 |
| Right medial posterior prefrontal cortex 1 | Control | 0.0146 | 1.16 | 0.0005 | 0.0037 |
| Left somatomotor 5 | Somatomotor | 0.0145 | 1.19 | 0.0005 | 0.0037 |
| Left precuneus 1 | Control | 0.0144 | 1.23 | 0.0014 | 0.0050 |
| Right dorsal attention precentral ventral 1 | Dorsal attention | 0.0141 | 1.09 | 0.0012 | 0.0048 |
| Left visual 8 | Visual | 0.0140 | 0.99 | 0.0028 | 0.0082 |
| Right temporo-occipito-parietal junction 2 | Salience / ventral attention | 0.0136 | 1.07 | 0.0007 | 0.0039 |

**Supplementary Table S7. Regions showing the largest between-treatment differences in vortex-space regional GBC.** The top 20% of Schaefer-100 regions are ranked by the absolute treatment difference, defined as mean ΔGBC following psilocybin minus mean ΔGBC following escitalopram, where ΔGBC denotes the post- minus pre-treatment change. Positive values thus indicate greater GBC changes following psilocybin relative to escitalopram.

| Group | Age | Gender | Ethnicity | Employment status |
|---|---|---|---|---|
| Psilocybin | 41.9 ± 11.0 years | 14 M<br>8 F | 14 White British<br>7 Other White<br>1 Mixed White/Asian & White British | 15 Employed<br>4 Unemployed<br>2 Student<br>1 Retired |
| Escitalopram | 38.7 ± 11.0 years | 14 M<br>6 F | 15 White British<br>2 Other White<br>1 Mixed Asian Indian + White British<br>1 Asian Indian<br>1 Pakistani British | 15 Employed<br>4 Unemployed<br>1 Student |

**Supplementary Table S8. Self-reported demographic characteristics of participants included in the study.** Age is reported as mean ± s.d.; all other values indicate participant counts.
